# Localized Space Coding and Phase Coding Complement Each Other to Achieve Robust and Efficient Spatial Representation

**DOI:** 10.1101/2025.09.07.674775

**Authors:** Tianhao Chu, Yuling Wu, Wentao Qiu, Zihao Jiang, Neil Burgess, Bo Hong, Si Wu

**Affiliations:** School of Psychological and Cognitive Sciences, Peking University, China; Peking-Tsinghua Center for Life Sciences, Academy for advanced interdisciplinary studies, Peking University, China; Beijing Key Laboratory of Behavior and Mental Health, IDG/McGovern Institute for Brain Research, Center of Quantitative Biology, Peking University, China; Department of Neurobiology, Northwestern University. Evanston, IL, USA; To Yuen Building, City University of Hong Kong, 31 To Yuen Street, Kowloon Tong, Kowloon, Hong Kong SAR; Institute of Cognitive Neuroscience, Department of Neuroscience, Physiology and Pharmacology, University College London, UK; Medical School Building B204 Tsinghua University Beijing, 100084, China

## Abstract

Localized space coding and phase coding are two distinct strategies responsible, respectively, for representing abstract structure and sensory observations in neural cognitive maps. In spatial representation, localized space coding is implemented by place cells in the hippocampus (HPC), while phase coding is implemented by grid cells in the medial entorhinal cortex (MEC). Both strategies have their own advantages and disadvantages, and neither of them meets the requirement of representing space robustly and efficiently in the brain. Here, we show that through reciprocal connections between HPC and MEC, place and grid cells can complement each other to overcome their respective shortcomings. Specifically, we build a coupled network model, in which a continuous attractor neural network (CANN) with position coordinate models place cells, while multiple CANNs with phase coordinates model grid cell modules with varying spacings. The reciprocal connections between place and grid cells encode the correlation prior between the sensory cues processed by HPC and MEC, respectively. Using this model, we show that: 1) place and grid cells interact to integrate sensory cues in a Bayesian manner; 2) place cells complement grid cells in coding accuracy by eliminating non-local errors of the latter; 3) grid cells complement place cells in coding efficiency by enlarging the number of environmental maps stored stably by the latter. We demonstrate that the coupled network model explains the seemingly contradictory experimental findings about the remapping phenomena of place cells when grid cells are either inactivated or depolarized. This study gives us insight into understanding how the brain employs collaborative localized and phase coding to realize both robust and efficient information representation.

## 1 Introduction

Place cells in the hippocampus (HPC) and grid cells in the medial entorhinal cortex (MEC) are two primary types of neurons involved in spatial representation [1, 2, 3, 4]. They adopt distinct strategies to encode space. Specifically, a place cell fires at a specific location in the environment, displaying a localized receptive field, referred to as localized space coding. While, a grid cell fires at multiple locations, displaying a periodic pattern across the environment (Figure 1a), and since all these periodic locations can be mapped into a single phase value in a period, this strategy is also called phase coding [5, 6, 7]. Interestingly, this dual coding scheme, i.e., the simultaneous use of localized and phase coding, is not restricted to spatial representation [8, 9]. Recent experimental studies have found that the similar scheme is applied in the representations of sound frequency [10] and gaze location [11]. For example, in the representation of sound frequency, both localized frequency coding and phase coding exist in the HPC and MEC, similar to how they are used in spatial representation [10, 8].

**Figure 1:**
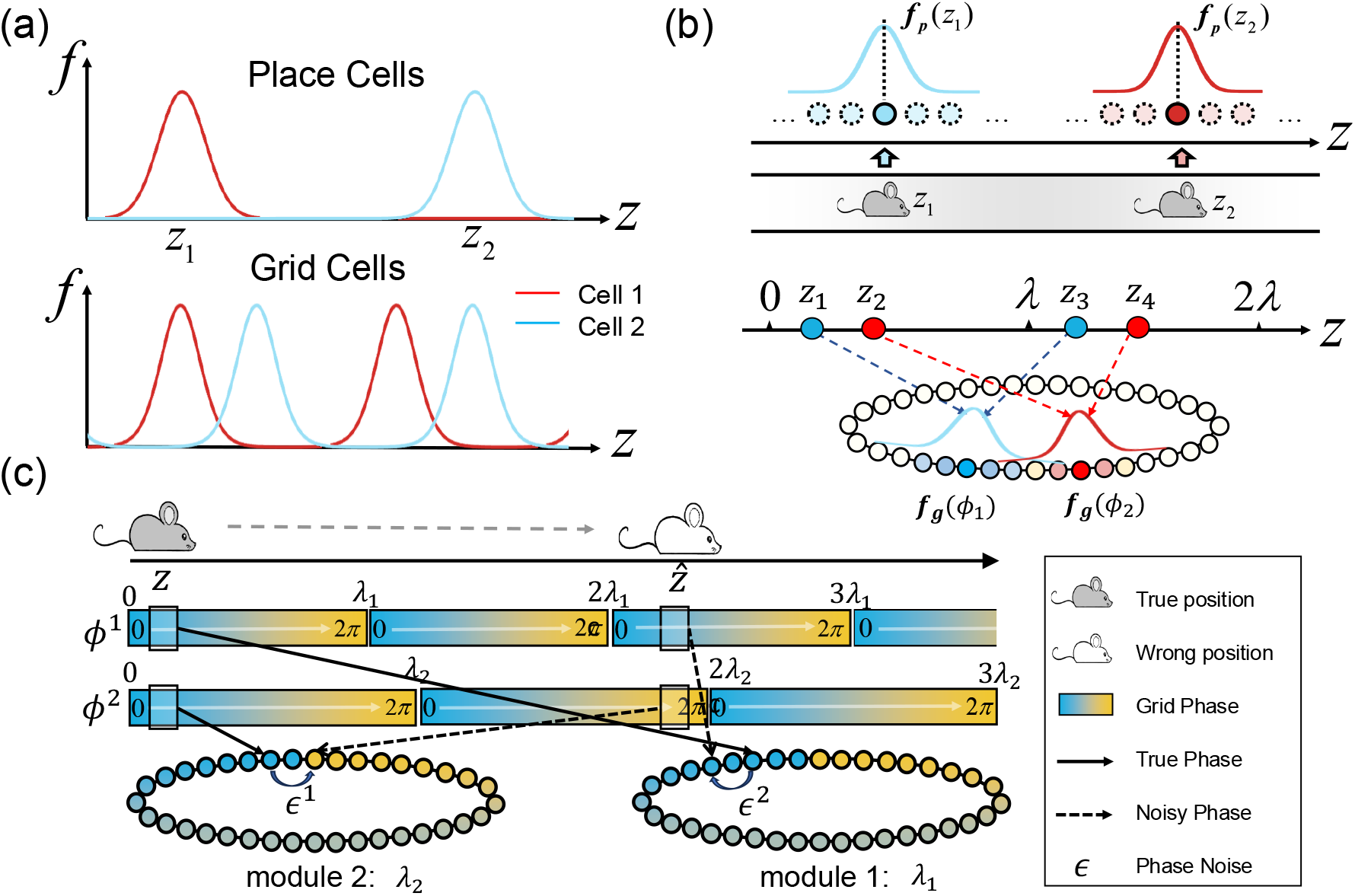
Illustrating coding properties of place and grid cells. **(a)**. Tuning curves of two example place cells and two example grid cells. A place cell fires at a single location (upper panel), while a grid cell fires at multiple periodical locations in the space (lower panel). **(b)**. Illustrating different coding strategies of place and grid cells. Upper panel: Local coding of place cells. The population activity of place cells forms a localized bump encoding the animal’s position. Lower panel: Phase coding of grid cells. Two equally spaced positions (*z*_1_, *z*_3_) or (*z*_2_, *z*_4_) with spacing *λ* are encoded by a single phase *ϕ*_1_ or *ϕ*_2_, according to the conversion rule *ϕ*(*z*) = mod(*z/λ*, 1) *×* 2*π − π*. **(c)**. Illustrating non-local errors of phase coding. An animal’s position *z* is represented by the combination of two phase values *ϕ*^1^ and *ϕ*^2^ from two grid cell modules with spacings *λ*_1_ and *λ*_2_, respectively. Small fluctuations in phases, denoted as *ϕ*^1^ + *ϵ*^1^ and *ϕ*^2^ + *ϵ*^2^, can incur a large error in spatial representation, from the true location *z* to a far away position 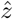.

Previous theoretical works have interpreted the dual coding scheme in the framework of cognitive map theory [8, 12, 9, 13]. According to this theory, there is a need for separate representations of abstract structure and sensory observations, and the dual coding scheme provides such computational support. Specifically, phase coding supports the representation of abstract transitions between concepts, while localized coding supports the association between abstract concepts and real-world observations [12]. In consistency with this view, it has been found that grid cells’ spatial representation is context-invariant and self-motion-based, reflecting the transition structure of the space; while place cells’ spatial representation is closely associated with external environments and exhibits remapping when the environment changes [14, 15, 16, 17]. The separation between abstract structure (phase coding) and sensory observations (localized coding) enables the neural system to generalize and reason in the conceptual space, a core idea of the cognitive map theory.

While the cognitive map theory and followed theoretical studies have highlighted the importance of the dual coding scheme, exactly how the two coding strategies work together to achieve optimal spatial representation remains largely unclear. In particular, it is known that either localized space or phase coding has its own distinctive shortcomings, and none of them is adequate to achieve robust and efficient space representation. Shortly speaking, localized coding, which utilizes a large number of neurons with overlapped local receptive fields to cover the feature space compactly (Figure 1a-b, upper panel), is robust to noise but not efficient, since the number of neurons it recruits increases linearly with the size of the environment [18]. On the other hand, phase coding, which utilizes the combination of grid cells with varying spacings to represent spatial location (Figure 1a-b, lower panel), is efficient but vulnerable to noise, since a small fluctuation in grid cells’ activity can lead to large, non-local errors in spatial representation [5, 6, 19] (Figure 1c). Given their respective advantages and disadvantages, it prompts a question: whether localized and phase coding can collaborate with each other to complement their shortcomings, such that the neural system achieves both robust and efficient information representation?

In this study, we use space representation as an example to explore how localized coding (place cells) and phase coding (grid cells) interact to complement each other’s shortcomings. Specifically, we construct a computational model mimicking the circuit of reciprocally connected HPC and MEC. The model consists of a number of continuous attractor neural networks (CANNs), where a CANN with position coordinate (P-CANN) represents the place cell ensemble, and a set of CANNs of phase coordinates (G-CANNs) represent grid cell modules of varying spacings. The P-CANN and each G-CANN are reciprocally connected in a congruent manner, conveying the correlation prior between the sensory cues processed by HPC and MEC, respectively. We show that within the framework of probabilistic inference, the dynamics of this coupled network effectively implement gradient-based Bayesian integration of sensory cues.

Through both theoretical analysis and simulation studies, we demonstrate that the coupled HPC-MEC network achieves complementary coding in three aspects: 1) place and grid cells interact to integrate sensory cues in a Bayesian manner; 2) place cells complement grid cells in coding accuracy by eliminating non-local errors of the latter; 3) grid cells complement place cells in coding efficiency by enlarging the number of environmental maps stored stably by the later. Furthermore, we show that the coupled network model provides a unified explanation for the seemingly contradictory experimental findings related to the remapping phenomena of place cells when grid cells are either inactivated or depolarized [20, 21]. We hope this study gives us insight into how localized and phase coding coordinate to achieve optimal representation in general cognitive maps.

## 2 Results

### 2.1 A network model with coupled place and grid cells

We build a network model mimicking the reciprocally connected HPC and MEC (see Figure 2a). A large volume of experimental and theoretical studies has suggested that CANNs are a canonical model for simulating spatial representation in the hippocampal-entorhinal system [22, 23, 24, 25, 26, 19, 27], we therefore adopt a CANN to model the sub-network formed by the place cell ensemble in HPC, referred to as P-CANN hereafter, and we adopt a set of CANNs, with each of them modeling the subnetwork formed by each grid cell module in MEC, referred to as G-CANNs hereafter. As indicated by experimental data, there is few connection between G-CANNs, while the P-CANN and each G-CANN are abundantly connected in a reciprocal manner [28]. For the clearance of description, we only present the results for the case of one-dimensional (1D) spatial representation, corresponding to the 1D linear track setting in experiments (e.g., [29]), but extension to the 2D case is straightforward.

**Figure 2:**
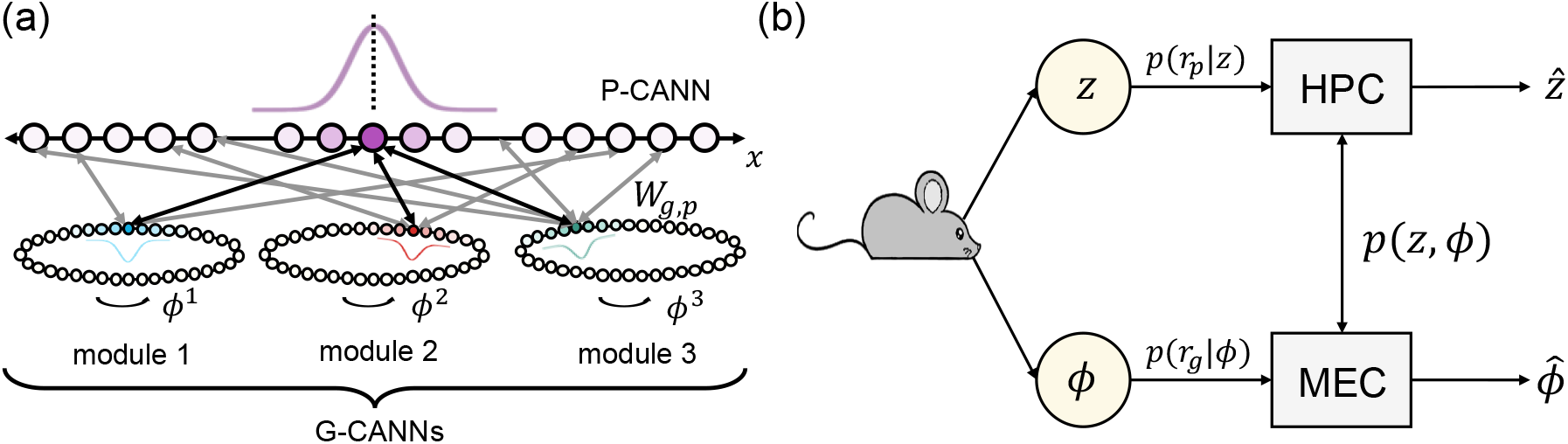
Complementary coding of space with place and grid cells. **(a)**. A network model with coupled place and grid cells. The place cell ensemble is modeled as a 1D CANN with location coordinate (P-CANN). Each grid cell module is modeled as a 1D CANN with phase coordinate (G-CANN). The P-CANN and each G-CANN are reciprocally connected, and there is no connection between G-CANNs. **(b)**. A probabilistic inference model of information integration between place and grid cells. The animal location *z* is encoded by the population activity **r**_*p*_ of place cells in HPC in the form of the likelihood function *p*(**r**_*p*_|*z*), and is represented by the population activity **r**_*g*_ of grid cells in MEC in the form of the likelihood function *p*(**r**_*p*_|***ϕ***). By integrating two sensory cues with the correlation prior *p*(*z*, ***ϕ***), HPC and MEC output the animal location 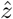 and the corresponding phase vector 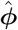, respectively.

Denote *z* ∈ (0, *L*) as the physical location of an animal. Neurons in the P-CANN are aligned according to their preferred locations *x* ∈ (0, *L*). To encode an animal location *z*, the P-CANN generates a localized activity bump centered at *z*. We consider *M* grid cell modules, and in each module, a grid cell responds to multiple periodic locations that are equally spaced by *λ*_*i*_, for *i* = 1, …, *M* . To uniquely quantify the receptive field of a grid cell, we define a phase value, which is given by

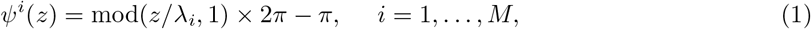

where the function mod(·) denotes the remainder operation, which converts the periodical locations *z* into a single phase value *ψ*^*i*^(*z*). In each G-CANN, grid cells are aligned according to their preferred phases *θ*^*i*^ ∈ (*− π, π*]. To encode an animal location *z*, each G-CANN generates a localized activity bump centered at *ϕ*^*i*^ = *ψ*^*i*^(*z*) in the phase coordinate.

Denote *U*_*p*_(*x, t*) and *U*_*g*_(*θ*^*i*^, *t*) the synaptic inputs, respectively, to place cells at location *x* in the P-CANN and grid cells at phase *θ*^*i*^ in the *i*th G-CANN, and *r*_*p*_(*x, t*), *r*_*g*_(*θ*^*i*^, *t*) the corresponding firing rates of neurons. The dynamics of the coupled network are expressed as,

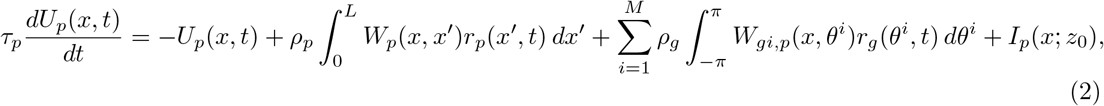

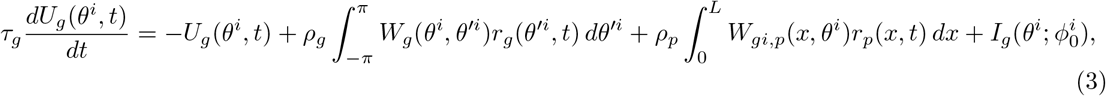

where *τ*_*p*_ and *τ*_*g*_ are synaptic time constants, and *ρ*_*p*_, *ρ*_*g*_ are neuronal densities. The other terms are introduced below.

#### Recurrent connectivity

*W*_*p*_(*x, x*^*′*^) and *W*_*g*_(*θ*^*i*^, *θ*^*′i*^) denote the recurrent connections between neurons in each CANN sub-network. They are translation-invariant in the feature space (*x* or *θ*) and decay with the difference between neuronal preferred locations or phases. They enable each CANN sub-network to hold a continuous family of bump states (see Methods Section 4.1 for detailed mathematical formulation).

#### Reciprocal connectivity

*W*_*gi*,*p*_(*x, θ*^*i*^) denote the reciprocal connections between place cells and grid cells. We set them to be congruent, in terms of that the connection strength is strong when the place cell’s location matches the grid cell’s phase according to the conversion rule (Eq (1)) (see Methods Section 4.1). This congruent form of connections imposes that the spatial representations of place and grid cells tend to agree with each other, reflecting the correlation prior between the environmental and self-motion cues.

#### Activation function

The neuronal firing rates *r*_*p*_ and *r*_*g*_ are determined by their synaptic inputs under the regularization of divisive normalization, which implicitly implements global inhibition, a mechanism widely observed in cortical circuits to regulate population activity [30] (see Methods Section 4.1 for mathematical formulation). It can be checked that without reciprocal connections, the attractor states of the P-CANN and each G-CANN are localized bump activities, which can be written 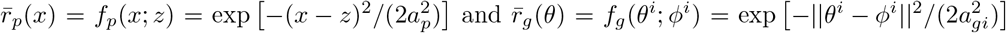, with *z* and *ϕ*^*i*^ representing the bump centers (see SI Section 3). If the network’s bump center perfectly represents the animal’s spatial location (in the absence of external noises), the steady-state functions *f*_*p*_(*x*; *z*) and *f*_*g*_(*θ*^*i*^; *ϕ*^*i*^) are the tuning curves of place and grid cells, respectively.

#### External input

Experimental studies have shown that place cells and grid cells are driven by environmental and self-motion cues, respectively [15, 16]. Accordingly, in the model we set the external input *I*_*p*_(*x*; *z*_0_) to place cells as a Gaussian bump centered on the animal’s true position *z*_0_ plus noises, representing the direct sensory observation of the animal location. On the other hand, for the external input 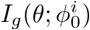 to grid cells, we calculate the phase 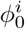 of grid cells through integrating the self-motion velocity of the animal and after that we set 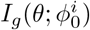 to be Gaussian bump centered at 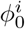 plus noises. The precise forms of *I*_*p*_(*x*; *z*_0_) and 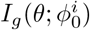 are given in Methods Section 4.1.

### 2.2 Bayesian interpretation of the coupled network dynamics

We first analyze that from the probabilistic inference point of view, the coupled network effectively implements Bayesian integration of the sensory cues received by place and grid cells.

#### 2.2.1 Low-dimensional dynamics of the coupled network

To analyze the computational process carried out by the coupled network, we first reduce the dimensionality of its dynamics. This utilizes the key property of a CANN, that is, the dynamics of a CANN is dominated by very few motion modes, in particular, the bump movement in the feature space [31, 32]. We can project the network dynamics on these dominating motion modes to reduce the complexity of the dynamics dramatically. After applying the projection method (see Methods Section 4.2), we obtain the dynamics of the activity bump center (*z*) in the P-CANN, which corresponds to the place cells’ representation of the animal location, and the dynamics of the activity bump center (*ϕ*^*i*^) in each G-CANN, which corresponds to the grid cells’ representation of the animal location (in the form of phase coordinate). They are written as,

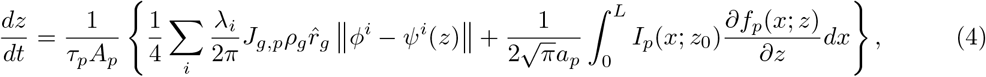

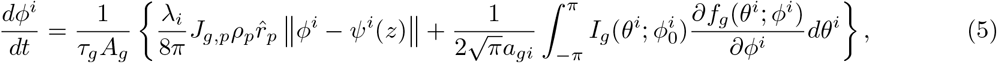

where 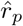 and 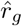 denote peak firing rates of place and grid cells, respectively. The first term on the right-hand side of each equation represents the contribution of the reciprocal interactions between place and grid cells, which tends to align the location representations of place and grid cells; the second term represents the contribution of the external input, which tends to align the neuronal representation with the sensory cue. The interplay between these two terms determines the network decoding result.

#### 2.2.2 The coupled network implements gradient-based Bayesian inference

We construct a probabilistic inference model to interpret the computation carried out by the coupled network (see Figure 2b).

#### Likelihood function

Place cells perceive the animal location through environmental cues, while grid cells perceive the animal location through the self-motion cue. Denote *p*(**r**_*p*_|*z*) the likelihood function of observing the joint activity **r**_*p*_ of place cells given the animal location *z*, and *p*(**r**_*g*_|***ϕ***) the likelihood function of observing the joint activity **r**_*g*_ of grid cells given the phase vector ***ϕ***(*z*) (which implicitly encodes the animal location *z*). Based on the theory of probabilistic population coding and the external input forms, we obtain the expressions of the likelihood functions *p*(**r**_*p*_ | *z*) and *p*(**r**_*g*_ | ***ϕ***) (see Method Section 4.3 and SI Section 2 for detailed mathematical form).

#### Correlation prior

Although the environmental and motion cues are processed through separate pathways in the brain, the spatial representation (*z*) of place cells and the phase representation (*ϕ*) of grid cells are not independent to each other, since they arise from the same signal resource (the true animal location). To account for this statistical correlation, we define the joint probability *p*(*z*, ***ϕ***) of observing the place cell bump center *z* and the phase vector ***ϕ*** = {*ϕ*^*i*^} of grid cell modules as:

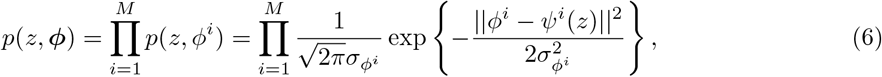

where *ψ*^*i*^(*z*) represents the converted phase of location *z* according to the conversion rule in Eq. (1). This Gaussian prior reflects that the animal’s locations estimated by place cells and grid cells tend to be aligned with each other, and the width 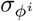 controls the correlation strength. Our studies show that this correlation prior can be realized by the congruent reciprocal connections between place cells and grid cells.

#### Bayesian inference

With the observed neuronal responses (**r**_*p*_, **r**_*g*_), the neural system infers the animal location *z* represented by place cells and the phase vector ***ϕ*** represented by grid cells. According to the Bayes’ theorem, the posterior distribution of (*z*, ***ϕ***) given (**r**_*p*_, **r**_*g*_) is written as

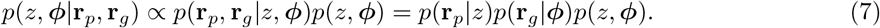

In the above, we have utilized the condition that the two sensory cues go through separate signal pathways. Mathematically, optimal decoding is achieved by maximizing the posterior (MAP). In practice, however, because of the high non-linearity of the posterior, it is often difficult to find a feasible algorithm to carry out the maximization operation, and approximation methods are often used. The gradient-based optimization of posterior (GOP) is such an approximation method [33]. It starts from an initial point and then ascends along the gradient of the log posterior to search for a local maximum solution. For the above probabilistic inference model, the gradients of the posterior are calculated to be (for details, see Methods Section 4.3 and SI Section 2),

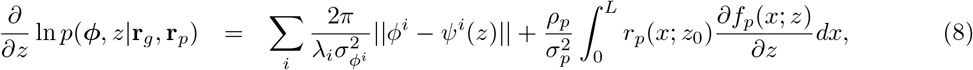

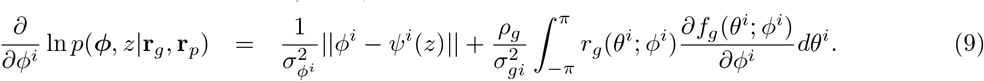

Comparing Eqs. (8-9) with Eqs. (4-5), we note that they have the same forms except for the coefficients. This implies that by setting the model parameters appropriately (see SI Section 3), the coupled network dynamics is equivalent to GOP. Interestingly, we find that in practice, because of the susceptibility of grid cells’ phases to noise, GOP can actually outperform MAP due to its history-dependent nature of computation (see below).

#### Numerical validation

We carry out numerical simulations to validate that the coupled network indeed realizes GOP. Consider a scenario, in which the animal’s true position is fixed at the center of a linear track of length *L* = 60, i.e., *z*_0_ = *L/*2. The network receives two noisy inputs, *I*_*p*_ and *I*_*g*_, representing the environmental and the self-motion cue, respectively. We decode the animal’s location and grid phases using two approaches: 1) GOP, searching the local maximum of the posterior along the gradient-ascent direction according to Eqs. (8-9); 2) Network decoding, evolving the coupled network dynamics according to Eqs. (2-3) to find the stationary states of the network.

Figure 3a displays examples of how the neural activity bumps in the P-CANN and G-CANNs move towards the animal location or the corresponding phase values under the drive of external inputs. Figure 3b shows that the trajectories of network decoding align with GOP very well. Figure 3c further shows that the results of network decoding agree well with GOP. We also validate the consistency between GOP and network decoding by varying model parameters (see SI Section 5). Overall, these numerical simulations confirm that the coupled network implements GOP effectively.

**Figure 3:**
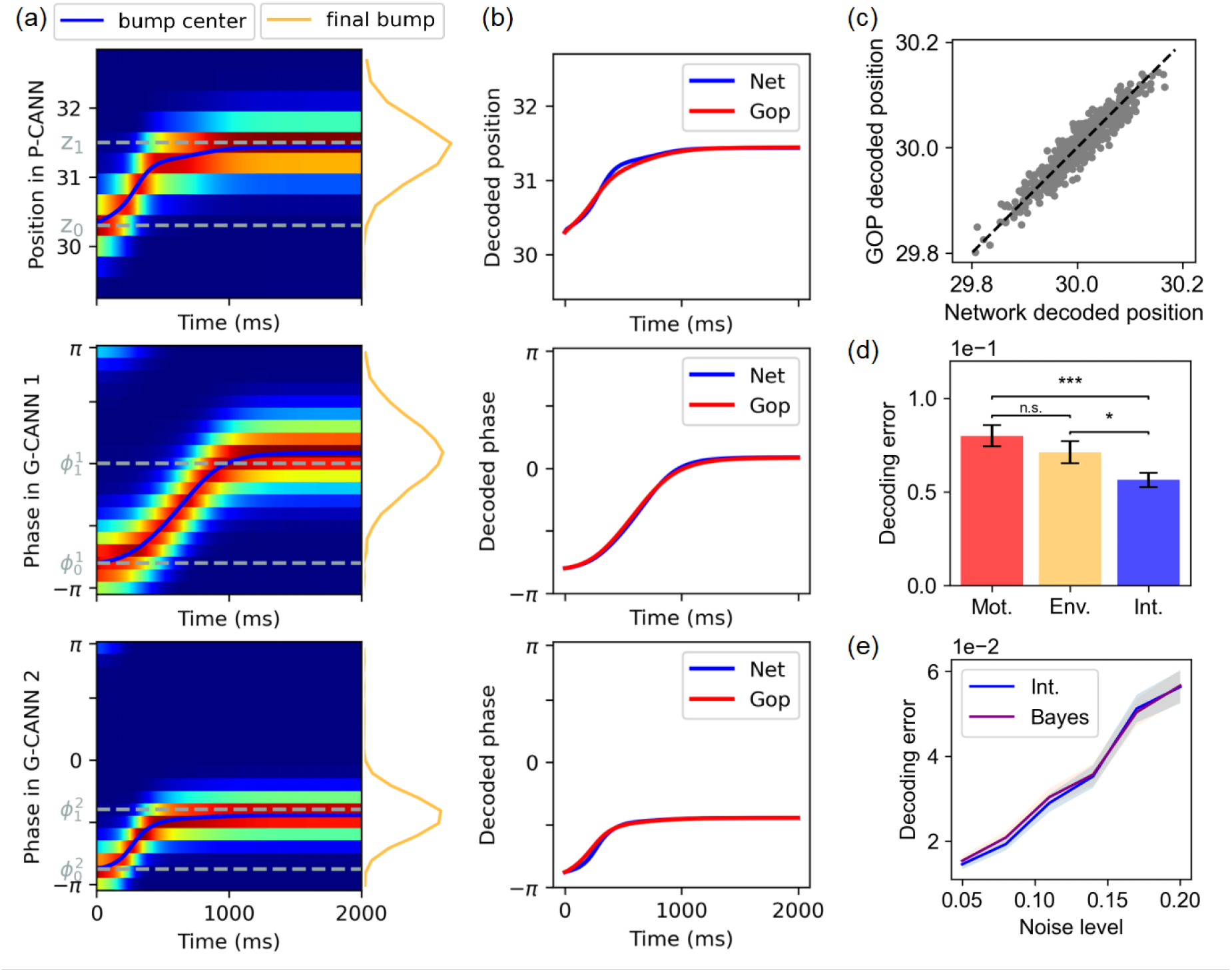
Bayesian interpretation of the coupled network dynamics. **(a)**. Illustrating the network decoding process. Neural activities in the P-CANN and two example G-CANNs are presented. The animal is initially located at *z*_0_ = 30.3, and the corresponding grid cell phases are 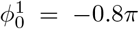 (G-CANN 1) and 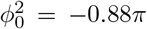 (G-CANN 2). The final animal location is at *z*_1_ = 31.5, and the corresponding grid cell phases are 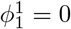 and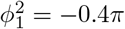 . Heatmaps showing neural activity dynamics over time versus decoded position and phase. Warmer colors represent higher firing rates. The solid blue lines indicate the temporal evolution of bump centers, while yellow lines at right show the final stabilized activity profiles. Under the drive of the external inputs, the bump center of the neural activity in the P-CANN moves towards the animal location *z*_1_ (upper panel), and the bump centers in two G-CANNs move towards the corresponding phase values 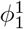 and 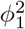, respectively (lower two panels). **(b)**. Comparing the trajectories of network decoding and GOP in the P-CANN (upper panel) and two G-CANNs (lower two panels). **(c)**. Network decoding vs. GOP. Each point represents the result of a single trial with different noisy inputs (500 trials). The dotted diagonal line indicates ideal perfect decoding. **(d)**. Network performances under different cuing conditions at noise level 0.2. Mot.: only the self-motion cue is presented (*I*_*g*_ ≠ 0, *I*_*p*_ = 0); Env.: only the environmental cue is presented (*I*_*g*_ = 0, *I*_*p*_ ≠ 0); Int.: both self-motion and environmental cues are presented (*I*_*g*_ ≠ 0, *I*_*p*_ ≠ 0). Error bars represent *±* SEM. Significance levels: ^*∗*^*p <* 0.05, ^*∗∗*^*p <* 0.01, ^*∗∗∗*^*p <* 0.001 (paired *t*-test, 100 trials). **(e)**. Network performance vs. Bayesian prediction under different noise levels. The network performance is measured by the averaged decoding error over 100 trials, shown as mean *±* SEM. The parameter used for simulations are given in Methods Section 4.1.2. Other simulation details are given in SI Section 4.

### 2.3 The coupled network achieving Bayesian information integration

We continue to demonstrate that the coupled network achieves Bayesian integration of sensory cues received respectively by place and grid cells. To this end, we consider three cuing conditions: 1) only the environmental cue is presented, corresponding to only place cells receiving external inputs (*I*_*p*_ ≠ 0 and *I*_*g*_ = 0); 2) only the motion cue is presented, corresponding to only grid cells receiving external inputs (*I*_*p*_ = 0 and *I*_*g*_ ≠0); 3) both the environmental and motion cues are presented, corresponding to both place and grid cells receiving external inputs (*I*_*p*_ ≠ 0 and *I*_*g*_ ≠ 0). In each condition, we run the network dynamics and read out the animal location by the bump position *z* of place cells. Figure 3d shows that the model integration yields lower decoding errors compared to that using a single cue. For comparison, we also calculate the theoretical prediction of Bayesian cue integration. Under the assumptions of Gaussian distributions of decoding errors and the uniform prior, the variance of Bayesian cue integration is given by 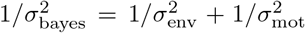, where 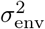 and 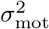 represent the variances of decoding errors of applying only the environmental or only the motion cue, respectively [34]. Figure 3e shows that the model decoding results agree well with the theoretical prediction of Bayesian integration across a wide range of noise levels, confirming that the coupled network integrates sensory cues optimally. For details of the experiments and calculations, see SI Section 4.

### 2.4 Complementary coding with coupled place and grid cells

Computationally, HPC implements localized coding via overlapping receptive fields of place cells, while MEC implements phase coding by combining the phases of grid cells of different spacings (Figure 1a–b). Previous studies have pointed out that localized coding is robust to noise but not efficient, since the number of neurons it utilizes increases linearly with the size of the environment [35, 18, 36, 37]; whereas, phase coding is efficient but not robust to noise, since small fluctuations in grid cells’ activities can incur large, non-local errors in spatial representation [5, 6, 19] (Figure 1c) (see more detailed analyses of the properties of localized and phase coding in SI Section 1). In the below, we show that through reciprocal connections, place and grid cells complement each other to overcome their respective shortcomings, and they work together to achieve both robust and efficient space representation.

#### 2.4.1 Complementary coding for robust space representation

We first demonstrate that place cells complement grid cells in coding accuracy. Let us consider a scenario, in which an animal is moving in darkness and only receives the motion cue (*I*_*p*_ = 0, *I*_*g*_ ≠ 0) (by only including the motion cue, it excludes the possibility that the improvement of the network performance is due to the integration of the environmental cue from place cells). We compare the network decoding results with and without reciprocal connections from place cells. In the later case, the animal position is decoded based solely on grid cells’ activities, i.e., by MAP based on grid cells’ activities (see simulation details in SI Section 4).

Figure 4a-c illustrate why MAP based on grid cells’ activities has large errors, whereas decoding by the coupled network has small errors. Figure 4d presents the one-step decoding results, which show that MAP based solely on grid cells’ activities can have very large errors when the noise strength is strong; whereas, these large errors are eliminated in the coupled network. Figure 4e further shows that as decoding errors are accumulated over time, even at a small noise level, the decoding error of MAP will become very large as time goes on; whereas, the decoding error of the coupled network remains to be constantly small.

**Figure 4:**
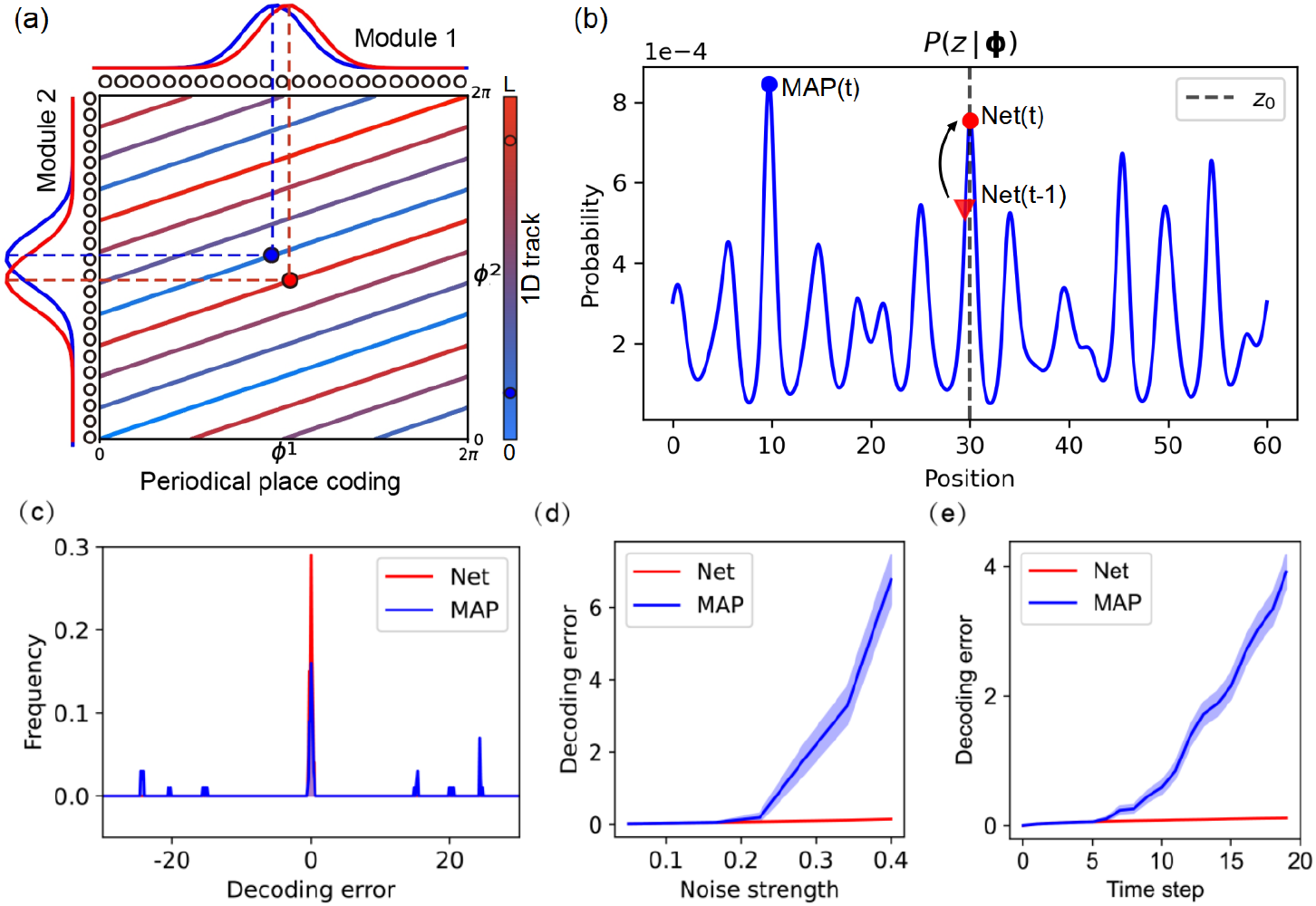
Place cells complement grid cells in coding accuracy. **(a)**. Illustrating the sensitivity of phase coding to noise. With two grid cell modules, a physical location (represented by color dots) is encoded as a unique combination of phases (*ϕ*^1^, *ϕ*^2^), corresponding to bump activities in two modules (shown at the top and left margins of the panel). Due to the periodic nature of grid cell phases, a linear track in the physical space is mapped into multiple parallel lines in the 2D phase space. As a result, two positions that are far apart in the physical space (red and blue dots) become close in the phase space. This proximity implies that small fluctuations in grid cell phases can cause large decoding errors. **(b)**. Illustrating large errors of MAP based on phase coding. The posterior distribution *P* (*z* | ***ϕ***) typically has multiple local maxima, and its global maximum (blue dot) may not align with the true animal location (*z*_0_) due to noise sensitivity, which causes large error of MAP. On the other hand, the decoding of the coupled network starts from the previous result (red triangle) and finds the local maximum (red dot). Because of the smooth movement of the animal, this leads to a robust decoding result. **(c)**. Example distributions of decoding errors (100 trials) of the coupled network (red line) and that of MAP based solely on grid cells’ activities (blue line). Note that MAP has high probability to generate large errors. **(d)**. One-step decoding results across different noise levels of the coupled network (red line) and MAP based solely on grid cells’ activities (blue line). **(e)**. Decoding results over time of path integration in the coupled network (red line) and MAP based solely on grid cells’ activity (blue line). Data is shown as mean *±* SEM across 200 trials. The parameter used for simulations are given in Methods Section 4.1.2. Other simulation details are given in SI Section 4.

The reason for that large errors are avoided in the coupled network can be intuitively understood as follows. As analyzed in Section 2.3, the dynamics of the coupled network effectively implement GOP, whereby the previous bump position of place cells (i.e., the previous estimated animal location) continuously serves as the starting point for the network to estimate the next-step animal position. This history-dependent way of computation naturally avoids abrupt change in decoding result, in consistent with the smooth movement of the animal. On the other hand, MAP based on grid cells’ activity alone calculates the global maximum of the posterior, and the latter is susceptive to noise (Figure 4b).

#### 2.4.2 Complementary coding for efficient space representation

While place cells form a robust localized representation of animal location in space, such a strategy is inefficient, since the number of neurons required to cover an environment grows linearly with the space size, restricting the coding efficiency of place cells [38]. Experimental studies have shown that the same population of place cells can be reused to encode multiple environments [39], often through remapping, whereby the receptive fields of cells change across environments. However, theoretical studies have long suggested that embedding multiple spatial maps within a single recurrent network leads to interference between stored patterns [40, 41, 25]. This interference arises because different attractor manifolds, each corresponding to a spatial map, rely on the same set of neurons, which causes overlap and instability in neural representation. Consequently, the number of distinct maps that can be reliably stored and retrieved by place cells, i.e., the representational capacity, is severely limited.

Here, we show that reciprocal connections from grid cells can mitigate such interference and substantially increase the representational capacity of place cells. This enhancement is due to two key properties of grid cells. First, grid cells provide a stable, environment-invariant metric representation of the space, undergoing only coherent realignments across environments [3]. Second, grid cells employ a combinatorial phase code across multiple modules, endowing them with an exponentially large representational capacity [5]. This high capacity of the coding space enables a crucial mechanism: when multiple spatial maps are embedded in the place cell network, their patterns can be projected into the grid cell space via reciprocal connections. In this higher dimensional space, these maps become more linearly separable and less likely to interfere. The grid cell representations, being stable and distinct, are then fed back to the place cell network, serving both as a reference metric and as a disambiguating scaffold that reinforces the separation between stored maps. As a result, the attractor dynamics of the place cell network become more robust, and the number of maps that can be reliably maintained is significantly increased.

To confirm this idea, we construct a multi-map model of P-CANN, in which multiple continuous attractor manifolds are embedded (Figure 5a). Specifically, the recurrent connectivity among place cells is set to be a normalized sum of *K* connection matrices, each encoding a spatial map (see Methods Section 4.4 for details). To evaluate the stability of each map, we initialize the network at an attractor state of a probe map *k*, denoted as 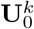, and then let the network evolve to a steady state, denoted as 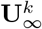 . The cosine similarity between 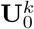 and 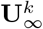, called bump score, measures the map stability.

**Figure 5:**
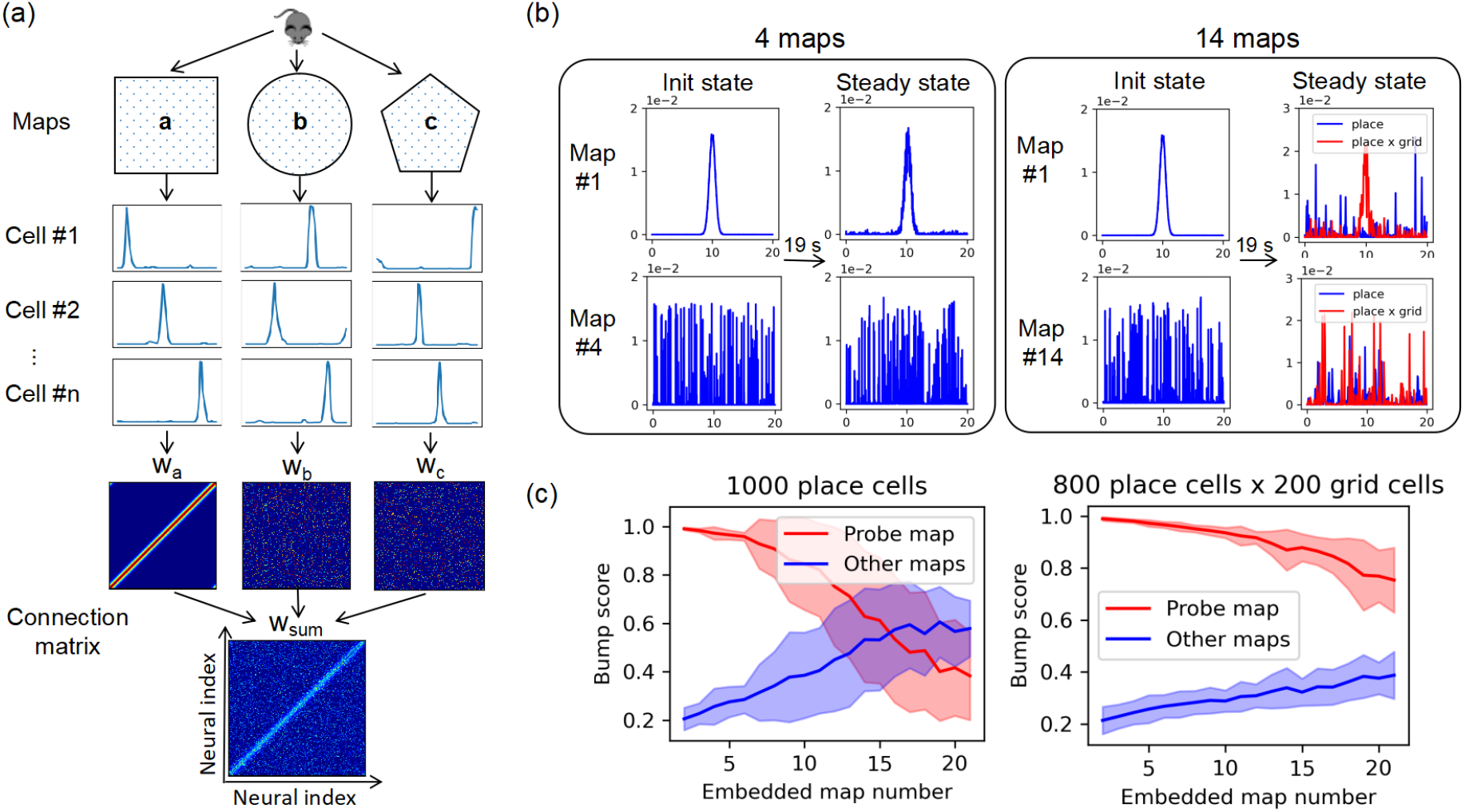
Grid cells complement place cells in coding efficiency. **(a)**. Illustration of the P-CANN encoding multiple environmental maps. Top panel: three environmental maps a, b, and c. Middle upper panel: neuronal receptive fields in different environments. Middle lower panel: neuronal recurrent connection matrices (*W*_*a*_, *W*_*b*_, *W*_*c*_) to support the corresponding maps. Note that neurons are aligned according to their preferred locations in map a, therefore only *W*_*a*_ exhibits the clear diagonal structure, while *W*_*b*_ and *W*_*c*_ do not due to the remapping. Bottom panel: the connection matrices for all individual maps are summed together with appropriate normalization, which define the connection matrix of the P-CANN (*W*_*sum*_). **(b-c)**. Grid cells enlarge the representation capacity of place cells. To compare fairly, the total number of neurons in the network is fixed (1000 place cells vs. 800 place cells + 200 grid cells). **(b)**. Illustration of stability/instability of an environmental map. Left panel: a map is stable in the P-CANN when four maps are encoded, even without reciprocal connections from grid cells. An initial attractor state of map 1 (upper column) is sustained after the network converging to the stationary state, whereas the state of map 4 (lower column) keeps random. Right panel: a map is unstable in the P-CANN alone when fourteen maps are encoded; while it becomes stable when the reciprocal connections between the P-CANN and G-CANNs are included. After the network converging to the stationary state, an initial attractor state of map 1 is changed (blue lines) or sustained (red line), depending on wether reciprocal connections from grid cells exist or not. **(c)**. The coupled network has a greater representation capacity than the P-CANN alone. As the number of embedded maps increases, the bump score (a measurement of stability) of the probe map decreases (red line), while the highest bump score of other maps increases (blue line). Left panel: without reciprocal connections from grid cells. Right panel: with reciprocal connections from grid cells. Data is shown as mean *±* SD across 100 trials. The parameter used for simulations are given in Methods Section 4.4. Other simulation details are given in SI Section 4.

We compare two conditions: one in which the P-CANN operates alone, and another in which the P-CANN receives reciprocal inputs from the coupled G-CANNs. Figure 5b presents examples, one in which the P-CANN alone can maintain a stable map, and the other in which the P-CANN alone cannot, but with reciprocal connections from grid cells, it can. Figure 5c-d show that as the number of stored maps increases, the bump score for the probe map declines due to the increased interference from other maps. However, the decline is consistently slower in the coupled network. We regard a map as being stable if its bump score surpasses that of other maps by a predefined margin (0.2 used in our simulations). Based on this criterion, we define the representational capacity as the maximum number of maps the P-CANN can store stably. As shown in Figure 5c-d, the coupled network achieves significantly higher representational capacity than the place-cell-only network (this phenomenon is robust with respect to parameter changes, see SI Section 5 for simulation results under other parameter regions).

The above results demonstrate that complementary coding with coupled place and grid cells not only contributes to increasing the robustness of position inference, but also enables a more efficient space coding by enlarging the number of encoded environments. This complementary division of coding strategy, that is, place cells encode specific environmental maps by using localized space code, while grid cells encode a global, high-capacity metric structure by using phase code, provides a principled solution to the trade-off between accuracy and efficiency in space representation [42].

### 2.5 Reconciling remapping phenomena of place cells

Experimental studies have reported seemingly contradictory findings on the role of MEC in place cell remapping. The experimental data in [20] showed that inactivating grid cells in MEC leads to place cell remapping; whereas the data in [21] showed that depolarization of grid cells induces place cell remapping. Here, we show that our model provides a unified explanation of these seemingly conflicting results.

To start, we first inspect how the reciprocal connection strength between grid and place cells (i.e., *J*_*g*,*p*_ in our model) affects the stability of a stored environment map. As shown in Figure 6a, the stability of a map depends on *J*_*g*,*p*_ in a non-monotonic way, i.e., the stability first increases with *J*_*g*,*p*_, reaching the maximum at 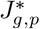, and then decreases. This result is intuitively understandable. When *J*_*g*,*p*_ is very small, increasing *J*_*g*,*p*_ implies enhancing the anchor effect of grid cells to place cells, which increases the stability of the map. However, when *J*_*g*,*p*_ is too large, the attractor dynamics of the place cell network is violated (since the recurrent inputs no longer dominate), and the network no longer performs as a CANN.

**Figure 6:**
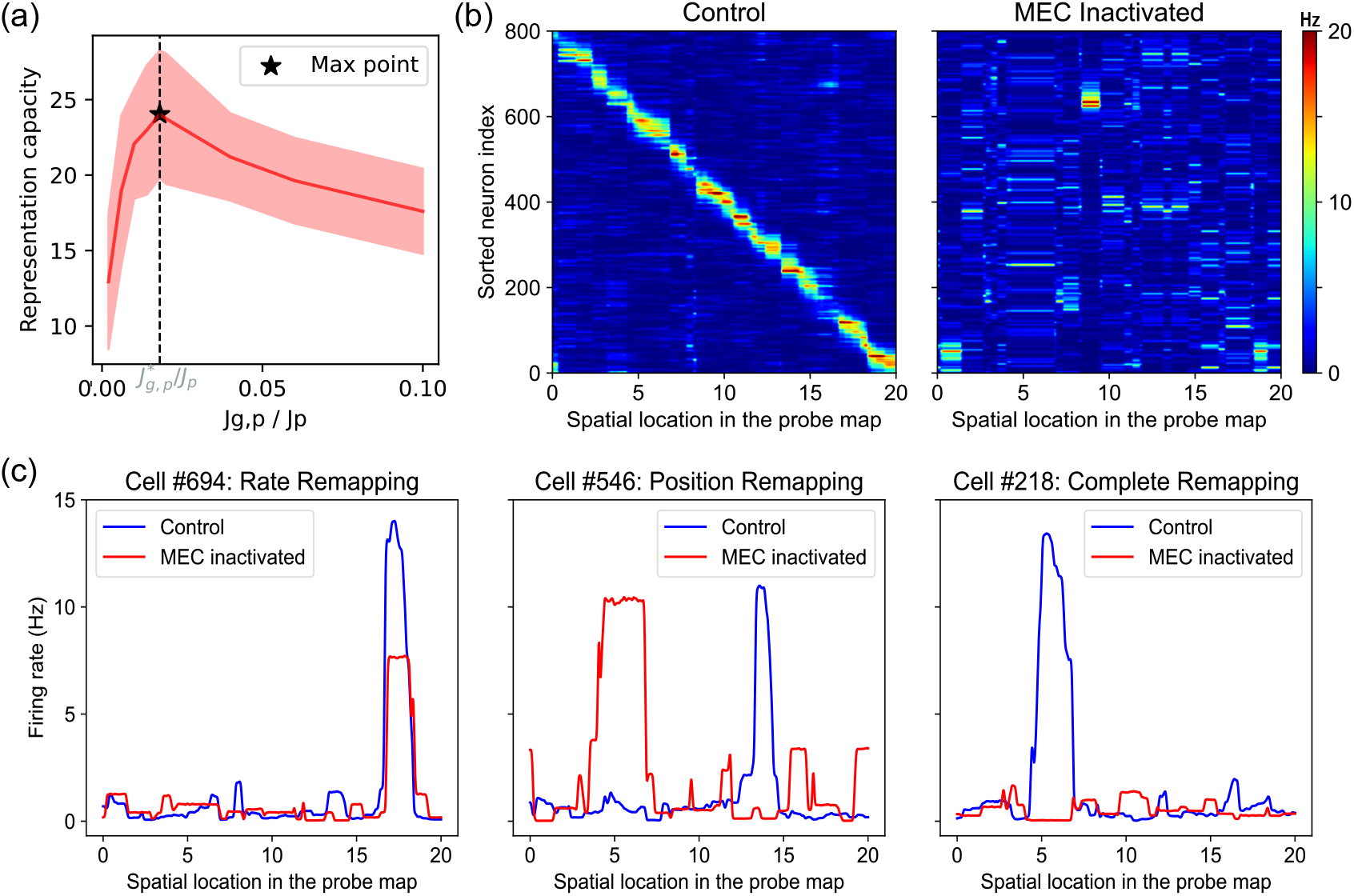
Inhibition of grid cells induces place cell remapping. **(a)**. The number of environment maps stored stably by the coupled network versus the ratio between the reciprocal connection strength *J*_*g*,*p*_ and the recurrent connection strength *J*_*p*_ of place cells, shown as mean ± SD across 30 trials. The dotted black line marks the optimal ratio 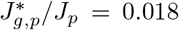 where the P-CANN has the maximum representation capacity. **(b)**. The remapped population activity of place cells. Left panel: the control condition. Right panel: MEC inactivated. **(c)**. Examples of three types of remapping at the single neuron level. Tuning curves of place cells under the control condition (blue line) and with inhibition of grid cells (red line). Left panel: rate remapping. Middle panel: place field remapping. Right panel: global remapping. The parameter used for simulations are given in Methods Section 4.5. Other simulation details are given in SI Section 4.

We can give an intuitive interpretation of the experimental data from the view of varying the effective value of *J*_*g*,*p*_ in our model. In the case of inactivating MEC (i.e., the input from grid to place cells becomes very weak), it is equivalent to setting *J*_*g*,*p*_ very small. As analyzed above, this reduces the stability of the environment map, resulting in that place cells’ activities shift to other maps due to the interference from other maps, namely, the remapping phenomenon occurs. On the other hand, depolarizing grid cells is equivalent to increasing the effective value of *J*_*g*,*p*_ while keeping activities of grid cells unchanged in our model. Consider that the practical value of *J*_*g*,*p*_ is a little larger than 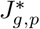 (the point having the maximum map stability). In such a case, increasing *J*_*g*,*p*_ leads to decreasing the stability of the environment map, which also induces the remapping of place cells.

We carry out simulations to formally reproduce the experimental data. Consider the case of inactivating MEC. We construct a coupled network model that can store *K* = 20 randomly generated environment maps stably. Consider that the network initially represents the probe environment map 1 (by setting the initial state of the network to be at an attractor state of the map 1). Mimicking the experimental protocol, we let an animal run through the environment *k* = 1 (see details in Methods Section 4.5) and measure the receptive fields of place cells as the animal running through. Two different parameter conditions are compared, *J*_*g*,*p*_ has the full value or a significantly reduced value (down to 0 in the simulation) . At the single neuron level, we obtain three remapping phenomena, which are: 1) rate remapping, where the place field of a neuron remains roughly unchanged, but its firing rate is altered (Figure 6c, left panel); 2) place field remapping, where the firing rate of a neuron remains relatively constant, but the location of the receptive field shifts (Figure 6c, middle panel); 3) global remapping, where the place cell of a neuron becomes inactive (Figure 6c, right panel) [43]. At the population level, we observe that the joint activity of place cells can no longer represent the original environment map faithfully (Figure 6b). Our model results agree very well with the experimental findings in [20]. The case of depolarizing of grid cells can be similarly studied and the results are presented in SI Section 4, which show that our model also reproduces the experimental findings in [21].

## 3 Conclusion and Discussion

In the present study, we have proposed that place and grid cells collaborate to complement their respective shortcomings, so that the brain achieves both robust and efficient representation of the space. Specifically, we built a computational model with reciprocally connected CANNs, in which one CANN with position coordinate models the place cell ensemble in HPC, a set of CANNs with phase coordinate model the grid cell modules of different spacing in MEC, and they are reciprocally connected in a congruent manner. We showed that this coupled network effectively implements the gradient-based Bayesian inference, achieving complementary coding in three aspects: 1) place and grid cells interact to achieve Bayesian integration of sensory cues they receive respectively; 2) place cells complement grid cells in coding accuracy by eliminating non-local errors of the latter; 3) grid cells complement place cells in coding efficiency by increasing the number of stable environments stored by the latter. We also demonstrated that this coupled network model can well explain the remapping phenomena of place cells when grid cells are either inactivated or depolarized.

### 3.1 Biological plausibility of the model

For the purpose of elucidating complementary coding clearly, we have adopted a simplified computational model and ignored many biological details. This simplified model, however, captures the main idea of space coding in the brain. A large number of experimental and theoretical studies have strongly suggested that CANNs are the suitable neural circuit model for describing spatial coding in HPC and MEC [22, 23, 24, 25, 26, 19, 27]. While place cells employ localized receptive fields [2, 44], grid cells employ periodic receptive fields to represent space [4], and they receive different types of sensory cues [16]. There are none or rather weak connection between grid cell modules [4], but there exist abundant reciprocal connections between HPC and MEC. In our model, we assume that the reciprocal connections between place and grid cells are congruent, in terms of that their connection strength is strong if the preferred location of the place cell matches the preferred phase of the grid cell. This is a biologically plausible assumption, since a pair of place and grid cells with matched preferred location and phase (i.e., they encode the same animal location) tend to fire together during animal navigation, and according to the Hebbian learning rule, their connection weight becomes strong. Nevertheless, in future work, we will extend the current model to include more biological details to unveil the detailed implementation of complementary space coding in the brain.

### 3.2 Biological implications of the model

The complementary coding theory helps us to understand two puzzles on space representation in the brain. The first one is about the spatial coding accuracy in darkness. In the dark condition, only grid cells receive the self-motion cue of the animal, and if only grid cells are involved to decode spatial location, the internal representation of animal’s location will become highly impaired, as phase coding has large non-local errors. However, through reciprocal connections with place cells (although place cells do not receive sensory cues in this case), the coupled network effectively implements GOP, which decodes the animal location based on the animal’s movement history and hence avoids large errors as demonstrated in our model.

The second puzzle is about the representational capacity of HPC. A well-known limitation of an attractor network is that its memory capacity is restricted due to the interference between memory patterns [41, 40, 25]. Our study shows that through reciprocal connections from grid cells (whose receptive fields are relatively stable with respect to environment changes), grid cells can serve as an anchor to stabilize the environmental map expressed in HPC. This effectively increases the number of stable environmental maps stored by the place cell ensemble. As an evidence of this anchor effect, when MEC is inactivated, the environment map hold by place cells loses its stability, and place cells exhibit the remapping phenomena, as demonstrated by both the experiment [20, 21] and our model (Figure 6). Notably, our model shows that the representation capacity of place cells increases with the number of grid cell modules, which is different to the result in [19]. We realize that this discrepancy comes from that the connection weights between place and grid cells in our model are normalized with respect to the number of grid cell modules. If no normalization is applied, we will get the same conclusion as in [19], since in such a case, as the number of grid cell modules increases, the reciprocal connections become very large, which destroy the attractor dynamics of the network, making our model no longer works.

### 3.3 Model predictions

In addition to reproducing the remapping phenomena of place cells as reported in the literature, our model also generates new predictions testable experimentally. Similar to inactivating MEC, one may inactivate HPC to induce changes in the activity pattern of grid cells and animal’s navigation behaviors. According to our model, after removing the inputs from place cells which convey the environmental cue information, the receptive fields of grid cells will become more stable with respect to environmental changes and exhibit reduced re-alignment (in our model, the re-alignment of grid cells is caused by the coupling with place cells). Moreover, as place cells no longer contribute to eliminating non-local errors of grid cells, we expect that the animal’s navigation becomes impaired. One may also inspect that whether the connection pattern between place and grid cells is congruent, a crucial assumption for the complementary coding theory. This could be done by measuring the connection strength or activity correlation between place and grid cells with matched/mismatched preferences. Furthermore, one can check the Bayesian integration between sensory cues. Current experiments using virtual reality can decouple spatial representation via motion or environmental cues, and show that spatial locations represented by place and grid cells can be different and they are influenced by both sensory cues, manifesting the information integration effect [45]. One may perform such experiment systematically to test whether or not the cue integration between place and grid cells is indeed Bayesian.

### 3.4 Comparison to previous works

The individual coding properties of place cells and grid cells have been intensively studied in the literature, including theoretically analyzing their respective advantages and disadvantages (see e.g. [46, 36, 35, 18, 5, 6, 47]). Recently, a number of modeling works have studied how place and grid cells interact with each other to support their respective functions. These include: 1) how the grid cell input drives place cells to form localized receptive fields and how the remapping phenomenon occurs [48, 19, 49]; 2) how the place cell input shapes the grid-like receptive fields of grid cells [50, 51, 52]; 3) how the HPC-MEC circuit supports multisensory processing. For instances, Li et al. proposed that reciprocal connections between HPC and MEC transmit the prior association between visual and motion cues, enabling the differentiation of multiple environments [53]; Laptev et al. suggested that the attractor dynamics in HPC and MEC enable decentralized information integration [15]; Agmon et al. modeled the joint network of grid cells and place cells as CANNs to perform spatial navigation [19]. In particular, Sreenivasan et al. pointed out the non-local error problem of grid cells and proposed a strategy to eliminate non-local errors by constraining the coding range of grid cells [6]. They also proposed a coupled network with connected place and grid cells to implement their strategy. However, in their model, place cells are not connected recurrently to form a CANN, and hence it is unable to realize GOP as in our model. Furthermore, the idea of “reducing the coding range” is at the cost of sacrificing the coding efficiency of grid cells, a key advantage of combinatory phase coding. Whereas, in our model, by considering the CANN dynamics of place cells, the coupled network not only reduces the coding sensitivity of grid cells but also retains their coding efficiency.

To our knowledge, our work is the first one that investigates the interaction between place and grid cells from the perspective of complementary coding. This theory is justified from both theoretical analysis and neural circuit implementation, which gives us insight into understanding how place and grid cells coordinate to improve both robustness and efficiency of spatial coding in the brain.

### 3.5 Generalization to general cognitive maps

In this study, we have focused on spatial representation. Previous works have suggested beyond spatial representation, the HPC-MEC formation may support general cognitive maps, where grid and place cells represent, respectively, abstract transition structure and associated sensory observations [13, 8, 12, 9]. Our model can be readily applied to these different cognitive maps, once abstract transitions can be formulated as “motion” in the phase space and sensory observations as “locations” in the feature space. This generality implies that the complementary coding framework implemented by interacted place and phase cells hold for general cognitive maps, and this dual coding scheme (localized and phase coding) serve as a principled strategy for the brain organizing maps of abstract knowledge [54].

### 3.6 Future Work

The present study has focused on investigating how place and grid cells complement each other to improve coding accuracy and efficiency. Experimental studies have suggested that place and grid cells may complement each other in other functions, such as in the construction of a global spatial map [49, 55]. Based on the environmental cues such as vision, place cells can perceive local space reliably but the view is restricted; wheres based on the self-motion cue, grid cells perceive space vaguely but can link far-away locations through path integration. Thus, to construct a global map, it need to combine information from both types of cues. In future research, we will explore how place and grid cells complement each other to construct a global spatial map.

## 4 Methods

### 4.1 The coupled network model embedded with a single spatial map

We model hippocampal place cells and entorhinal grid cells as two reciprocally connected continuous-attractor networks. The place-cell network (P-CANN) consists of *N*_*p*_ units on a 1D line *x* ∈ [0, *L*), and each of the *M* grid-cell modules (G-CANNs) consists of *N*_*g*_ units on a phase ring *θ* ∈ [*−π, π*).

The synaptic inputs *U*_*p*_(*x, t*) and *U*_*g*_(*θ*^*i*^, *t*) evolve according to

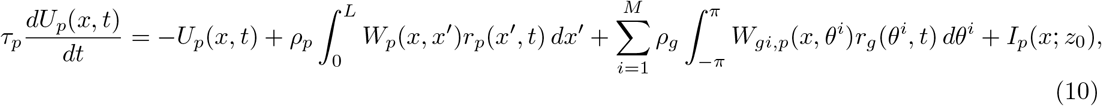

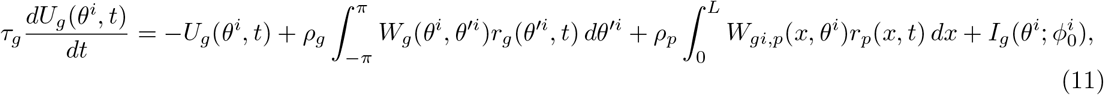

Recurrent weights of each CANNs are Gaussian kernels:

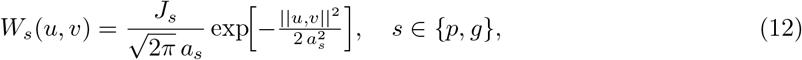

where ||*u, v* || is Euclidean for place cells with periodic boundary conditions to avoid edge effects, and circular distance for grid modules.

The reciprocal connections between place and grid cells are given by,

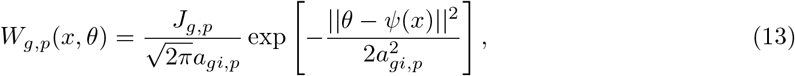

where *ψ*(*x*) = mod(*x/λ*, 1) × 2*π − π*.

Firing rates use divisive normalization:

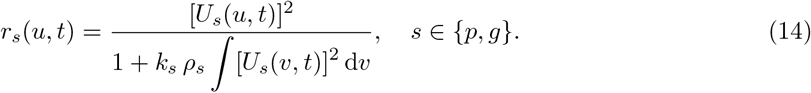

External drives reflect distinct sensory pathways. The place-cell network receives environmental observations directly from the animal’s true position:

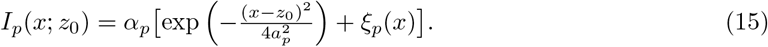

Grid cell modules implement path integration internally, a mechanism widely studied in prior work [26, 6]. Since our focus is on place–grid interactions rather than integration algorithms, we adopt a simple explicit update: at timestep *n*, module-*i* path-integrated phase 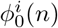 is

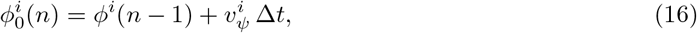

where *ϕ*^*i*^(*n −* 1) is the phase decoded from grid activity in the previous step and 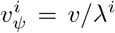 is the velocity in phase space. The external input to the grid module *i* is then

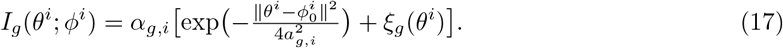

Decoding of *ϕ*^*i*^(*t*) from grid cell population activity and of position *z* from place cell activity will be detailed below.

#### 4.1.1 Bump-center decoding of *z* and *ϕ*^*i*^

It is well known that CANN can hold a continuous family of attractor states called “bump” states [56, 57, 58]. We mathematically prove that, under certain parameter constraints, P-CANN and G-CANNs can jointly support a continuous family of attractor states, among which each attractor state is supported by a set of coordinated bumps in P-CANN and G-CANNs (see SI Section 3).

We decode the represented position *z*(*t*) and phase *ϕ*^*i*^(*t*) by computing the center of the activity bumps of P-CANN and each G-CANN via complex-plane averaging on the ring:

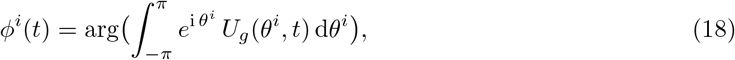

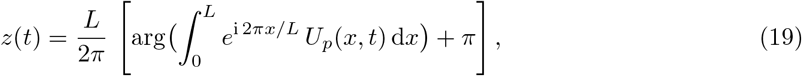

where arg() returns the complex argument in (*− π, π*], ensuring correct handling of periodic boundary of the network activities.

#### 4.1.2 Model Parameters

In our simulations, which demonstrate the position decoding accuracy in a single map (including the results shown in Figure 3 and Figure 4), we consider a 1D spatial range of *L* = 60. The model parameters include the following:

- **P-CANN parameters**: the connection width *a*_*p*_, the connection strength *J*_*p*_, the global inhibition strength *k*_*p*_, and the time constant *τ*_*p*_.
- **G-CANN parameters**: there are three grid cell modules in total. The parameters of each module *i* includes: the connection width for each module *a*_*gi*_, the connection strength between grid cells *J*_*g*_, the connection strength between P-CANN and G-CANNs *J*_*g*,*p*_, the global inhibition strength for each grid module *k*_*gi*_, and the time constant for each G-CANN *τ*_*gi*_.
- **Network structure parameters**: the number of place cell neurons *N*_*p*_, the number of grid cell neurons within each module *N*_*g*_, and the number of grid cell modules *M* .
- **Spatial properties**: the spacing of each G-CANN *λ*_*i*_.

The values of these parameters are listed in Table 1.

**Table 1:**
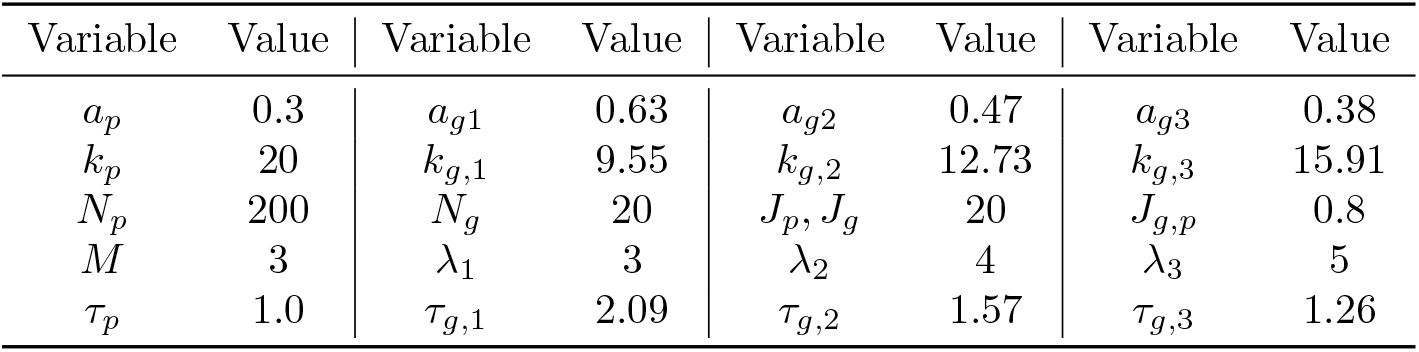
Parameters of the coupled network encoding a single map.

### 4.2 Simplifying the model dynamics by the projection method

The motivation behind simplifying the network dynamics lies in the need to study how the bump center of the network, which represents the position *z* and phase *ϕ* encoded by place and grid cells, evolves with noisy external inputs.

To achieve this, we use a projection method to reduce the high-dimensional neural dynamics of the network into the low-dimensional dynamics of the bump centers. This simplification not only makes the analysis more tractable but also reveals that the evolution of the bump center is consistent with the Gradient-based Optimization of Posterior (GOP) approach used as a way of Bayesian inference (detailed below). Here we outline how this simplification is carried out.

In a CANN, the stationary states form a neutrally stable sub-manifold. This suggests that the network dynamics can be effectively represented by a small number of dominant motion modes, such as variations in the height and position of these stationary states. Projecting the network dynamics onto these motion modes means multiplying the network dynamics by the corresponding motion mode and integrating over the relevant variable (either *x* for place cells or *θ* for grid cells). This projection greatly reduces the complexity of the system.

For place cells, the first two dominant motion modes are defined as the “height” mode 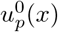 and the “position” mode 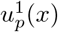, which are given by:

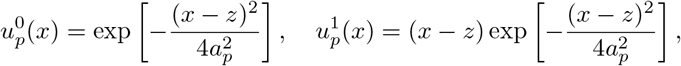

where *a*_*p*_ is the width of the place field, and *z* represents the position of the bump center for the place cells.

For grid cells, the dominant modes are similarly defined in phase space, with the height and position as:

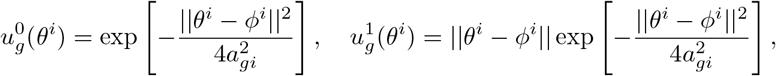

where *a*_*gi*_ is the grid spacing for module *i*, and *ϕ*^*i*^ represents the phase of the grid cell module.

The network dynamics are then simplified by projecting onto these modes. For place cells, the left-hand and right-hand sides of the height dynamics (projecting the network dynamics onto “height” mode) can be simplified to:

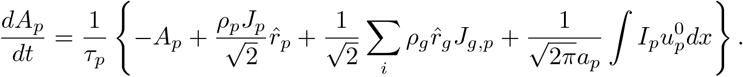

and the position dynamics (projecting the network dynamics onto “position” mode) are given by:

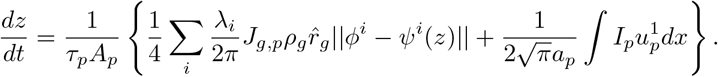

Similarly, for grid cells, the simplified projection leads to the following dynamics for phase:

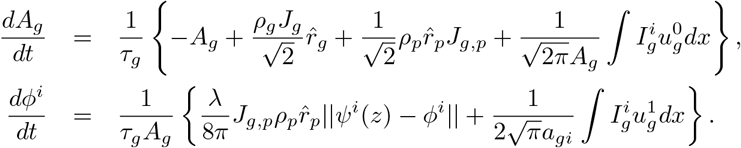

These simplified dynamics allow for more efficient simulation and analysis of the network’s behavior, while also providing insights into how the bump centers evolve under external noise and inputs.

For full details on the derivations and the final expressions, see SI Section 3.3.

### 4.3 Bayesian inference probabilistic model

Here, we introduce the details of the probabilistic inference model which is used for understanding the dynamical behavior or the network dynamics.

#### The encoding process

To reflect the correlation between the space representation of place cells and grid cells, we set the joint probability *p*(*z*, ***ϕ***) of the perceived animal location *z* by place cells and the estimated phases ***ϕ*** by grid cells satisfy,

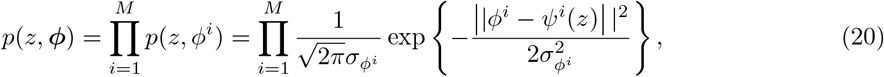

where *ψ*^*i*^(*z*) is the phase value matching the location *z*, i.e., *ψ*^*i*^(*z*) = mod(*z/λ*_*i*_, 1) *×* 2*π*. This Gaussian probability reflects that the animal location information perceived by place cells and grid cells tend to match each other statistically, and the Gaussian width 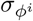 controls the correlation level between two sensory cues.

Without loss of generality, we consider Gaussian noises. The likelihood function of observing the joint activity **r**_*p*_ of the place cell ensemble given the animal location *z* is written as,

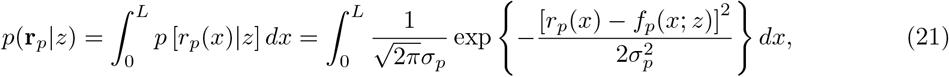

and the likelihood function of observing the joint activity **r**_*g*_ of grid cells given the phases ***ϕ*** is written as,

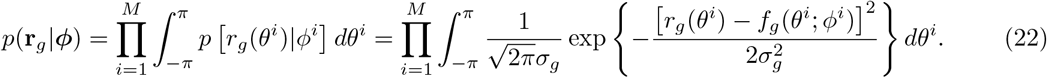

#### The decoding process

Given the observed responses (**r**_*p*_, **r**_*g*_) of place and grid cells, the neural decoder infers the animal location *z* represented by place cells and the phases ***ϕ*** represented by grid cells. According to the Bayes’ theorem, the posterior distribution of (*z*, ***ϕ***) given (**r**_*p*_, **r**_*g*_) is written as

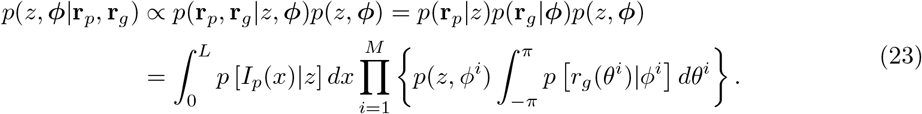

We consider the gradient-based optimization of posterior (GOP). Combining Eqs. (6-23), the gradients of the posterior are calculated to be (for details, see SI Section 2),

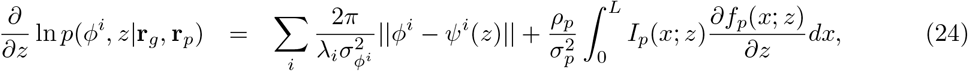

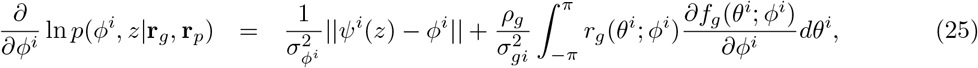

which give Eqs. (8,9) in the main text.

### 4.4 Embedding multiple maps in the coupled network

We construct the place cell network P-CANN to hold multiple CANN manifold, each of which corresponds to one environment map. For the convenience of description, let us denote *z* the retinatopic coordinate of a place cell and *x*^*k*^ the preferred location of the place cell in the *k*-th map, for *k* = 1, …, *K*, with *K* the total number of maps stored in HPC (note that there is no retina-topic map in HPC, i.e., *x*^*k*^ ≠ *z* in general). Without loss of generality, we set *x*^1^ = *z*, and for *k* ≠ 1, we randomly shuffle the retina-topic coordinates of all neurons to get *x*^*k*^. This mimics the remapping of receptive fields of place cells when the environment changes. The recurrent connections between place cells at retina-topic locations *z* and *z*^*′*^ are set to be,

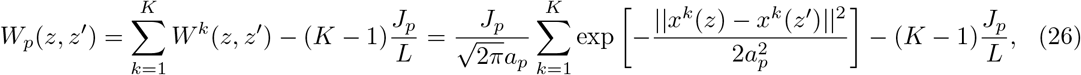

where *x*^*k*^(*z*) denotes the preferred location of the neuron at the retina-topic position *z*. Note that *W*^*k*^(*z, z*^*′*^) is the connection matrix that enables the place cell ensemble to hold a CANN in the feature space *x*^*k*^ (see Eq. 12), and *J*_*p*_*/L* is the mean of connection strength over all neuron pairs of each single map. Substracting this mean value avoids the connection strength monotonically increasing with the environment map.

The reciprocal connections between place cells and grid cells are adjusted accordingly, which are given by,

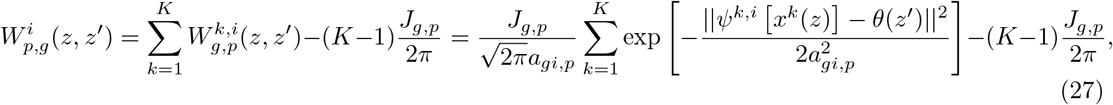

where 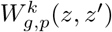 represents the connection strength between the place cell at the retina-topic location *z* in the *k*th map and the grid cell at the retina-topic location *z*^*′*^ in the *i*th module. *ψ*^*k*,*i*^(*x*^*k*^) = mod(*x*^*k*^*/λ*_*i*_, 1) *×* 2*π* + Δ^*k*,*i*^ converts position to phase, where Δ^*k*,*i*^ represents a random offset which is uniformly sampled from the range (0, 2*π*). *J*_*g*,*p*_*/*2*π* denotes the average of 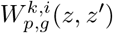 over all neuron pairs of each single map.

#### 4.4.1 Representation capacity of place cells

The representation capacity refers to the number of environment maps that can be stored stably by place cells. We differentiate between two conditions: with or without coupling from grid cells.

##### Bump Score Calculation

To evaluate whether the place cell network exhibits a bump state and to assess its representation of position *x* in a specific map *l*, we calculate the cosine similarity *q*^*l*^(*x*) between the actual population activity pattern of place cells and the idealized bump-shaped activity pattern corresponding to position *z* in map *l*:

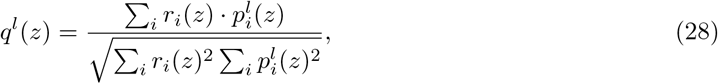

where *r*_*i*_(*z*) is the actual firing rate of place cell *i* at position *z*, and 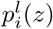 is the idealized Gaussian-curve firing rate of neuron *i* position *z* in map *l*. Next, we define a bump score for each map as the maximum cosine similarity 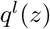 across all positions *z* in map *l*:

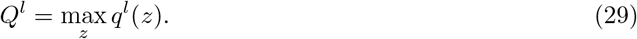

The map with the highest *Q*^*l*^ value is the winning map, and the position *z* corresponding to this value is the bump location for that map.

##### Simulation settings

The length of the environment is set to be *L* = 20. Without loss of generality, we consider the stability of the probe map *k* = 1. We set the initial state of the network to be at an attractor state of the network at the map *k* = 1. We let the network evolve for 1, 000 time steps with input and 19, 000 time steps without external input to obtain its stationary state. For each number of stored maps, we randomly generate networks with different permutations of place field orders and compute the mean and variance of the bump scores of the probe environment and the highest bump scores among the other environments.

##### Representation Capacity Calculation

We define that the network can no longer stably represent the probe map when the difference between the bump score of the probe map and the highest bump score of other environments fall below 0.2. This criterion allows us to determine the upper limit on the number of maps the network can encode, which corresponds to the network’s spatial representation capacity.

The default parameters of the coupled network is shown in Table 2.

**Table 2:**
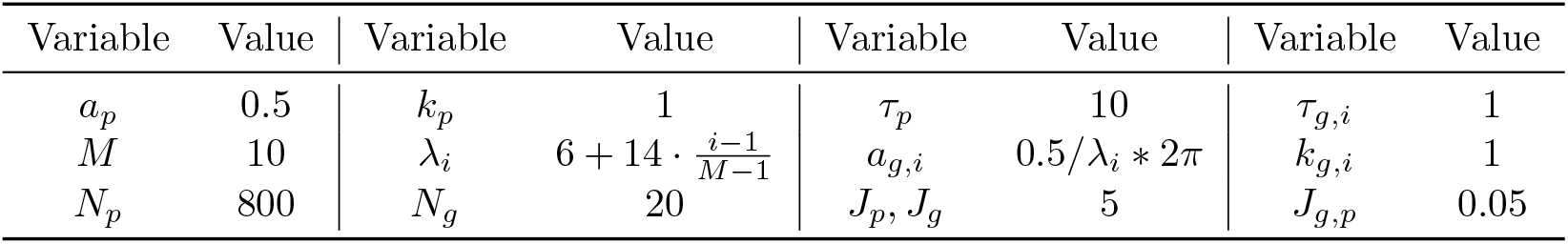
Parameters of the coupled network encoding multiple maps.

| Variable | Value | Variable | Value | Variable | Value | Variable | Value |
| --- | --- | --- | --- | --- | --- | --- | --- |
| $a_p$ | 0.5 | $k_p$ | 1 | $\tau_p$ | 10 | $\tau_{g,i}$ | 1 |
| $M$ | 10 | $\lambda_i$ | $6 + 14 \cdot \frac{i-1}{M-1}$ | $a_{g,i}$ | $0.5/\lambda_i * 2\pi$ | $k_{g,i}$ | 1 |
| $N_p$ | 800 | $N_g$ | 20 | $J_p, J_g$ | 5 | $J_{g,p}$ | 0.05 |

### 4.5 Replicating experimental findings on remapping phenomena of place cells

We introduce how we conducted simulation experiments to replicate two seemingly contradictory experimental observations regarding place cell remapping: one in which inhibiting MEC neuron activity induces remapping, and the other in which depolarizing MEC neurons induces remapping.

The coupled network was initialized to store 20 distinct environment maps. We simulated network dynamics by sampling 1000 uniformly distributed positions across the environment (length *L* = 20). For each position *z*_*i*_ (*i* = 1, …, 100), both grid and place cell activities were initialized to a bump state centered at *z*_*i*_, with external inputs applied for the first 100 ms to establish the initial activity pattern, followed by 19, 000 s of autonomous network evolution until reaching steady state. We compared two experimental conditions: (1) normal coupled network dynamics (control, *J*_*g*,*p*_ = 0.05 as the baseline coupling strength), and (2) manipulated grid-to-place cell projections through either complete inhibition (*J*_*g*,*p*_ = 0) or depolarization (*J*_*g*,*p*_ = 1). Final place cell activity patterns were recorded across all positions to construct complete spatial representations of each condition.

The simulation results for the inhibited grid cell condition are shown in Figure 6 in the main text, while the results for the stimulated grid cell condition are presented in SI Section 4.

Together, these simulation experiments demonstrate that our coupled network model can reproduce both the remapping induced by MEC inhibition and MEC depolarization, thereby providing a unified explanation for these seemingly contradictory experimental phenomena.

