## Supplementary information for "Localized Space Coding and Phase Coding Complement Each Other to Achieve Robust and Efficient Spatial Representation"

September 7, 2025

#### 1 Coding properties of place and grid cells

We review the different coding properties of place cells and grid cells. Local spatial coding (Localized Coding) by place cells, also known as "classical population coding" [1], has been widely studied in the literature (see e.g., [2; 3; 4; 5]). In this coding framework, each neuron has a local receptive field, which typically has the bell shape with its activity peaking at the center of the receptive field. Receptive fields of different neurons cover different parts of the space, together spanning the whole range. Place cells perceive the animal location primarily through environmental cues, e.g., the visual and/or olfactory cues. Denote the true location of the animal as $z$. The response of a place cell with preferred location at $x$ is written as,

$$r_p(x; z) = f_p(x; z) + \sigma_p \epsilon_p(x) = A_p \exp\left[-\frac{(x-z)^2}{4a_p^2}\right] + \sigma_p \epsilon_p(x), \tag{1}$$

where $f_p(x; z)$ is the tuning function, with $A_p$ and $a_p$ the peak and the width of the tuning function, respectively. $\epsilon_p(x)$ denotes independent Gaussian noise of zero mean and unit variance, and $\sigma_p$ the noise strength. In Localized Coding, a location is represented by the activity of a population of place cells, which forms a bump state of the network (Figure S1b).

Unlike place cells, grid cells respond at multiple spatial locations with periodicity. This can be equivalently described as grid cells have preferred phases in a period. In the 1D space, for grid cells with spacing $\lambda$, the match between phase and position is determined by a conversion rule [6],

$$\phi(z) = \mathrm{mod}(z/\lambda, 1) \times 2\pi - \pi, \tag{2}$$

where $z$ denotes the spatial position, and $\phi$ the corresponding phase value, ranging from 0 to $2\pi$. The operation mod denotes the modulus function. Grid cells perceive the animal location primarily through the self-motion cue. Denote $\phi^i(z)$ the phase of the $i$th module corresponding to the animal location $z$. The response of a grid cell in the $i$th module with preferred phase $\theta^i$ is written as,

$$r_g(\theta^i; \phi^i) = f_g(\theta^i; \phi^i) + \sigma_{gi}\epsilon_g(\theta^i) = A_g \exp\left\{-\frac{||\theta^i - \phi^i||^2}{4a_{gi}^2}\right\} + \sigma_{gi}\epsilon_g(\theta^i), \tag{3}$$

where $f_g(\theta^i; \phi^i)$ is the tuning function, with $A_g$ and $a_{gi}$ the peak and width of the tuning function, respectively. $\epsilon_g(\theta^i)$ represents Gaussian noise of zero mean and unit variance, and $\sigma_{gi}$ the noise strength. The operation $||\theta^i - \phi^i|| = \min\left(|\theta^i - \phi^i|, 2\pi - |\theta^i - \phi^i|\right)$.

Phase Coding employs a number of grid cell modules with different spacings to encode a spatial location [7], and they collectively form a phase vector (Figure S1c).

##### 1.1 Efficiency and robustness of Localized Coding

Previous studies have analytically derived the optimal decoding error of Localized Coding by calculating the Fisher information [8; 9; 10; 11]. These results show that the lower bound of decoding error of Localized Coding is linearly proportional to the noise intensity and inversely proportional to the total number of place cells. Specifically, under the assumption of independent Gaussian noises (see Eq. 1) and considering $x \in (-\infty, \infty)$, the Fisher information is calculated to be,

$$F(z) = \rho_p \frac{\sqrt{\pi} A_p^2}{2 a_p \sigma_p^2}, \tag{4}$$

where $\rho_p$ represents the density of place cells. We see that the Fisher information increases with the neuron density, indicating that increasing the number of neurons improves the robustness of Localized Coding.

In Localized Coding, the trade-off of increased robustness is the decrease of coding efficiency, as more neurons are needed to reduce the decoding error. Based on the information theory [12], Sreenivasan et al. defined coding efficiency $R$ as the total amount of encoded information divided by the number of encoding neurons [1]. For the 1D case, the total encoded information is proportional to the logarithm of the encoded spatial range $L$, so the coding efficiency is calculated to be $R = \ln L/N$. With this definition, we can roughly estimate the efficiency of Localized Coding in the 1D space. Assuming that place cells' place fields uniformly cover a 1D space, denoting $\Delta x$ to be the interval covered by a single place cell, the total range covered by $N$ neurons is $L = N\Delta x$. Therefore, the coding efficiency is calculated to be,

$$e_p = \frac{\ln(N\Delta x)}{N}. \tag{5}$$

As we can see, as the number of neurons $N$ increases, the efficiency $e_p$ of Localized Coding decreases and asymptotically approaches zero, indicating that Localized Coding is not efficient in terms of utilizing the encoding resource of neurons.

#### 1.2 Efficiency and robustness of Phase Coding

In Phase Coding, a single phase can only represent a position unambiguously within the range $\lambda$, beyond this range, ambiguity arises due to the periodical responses of grid cells. To enlarge the coding range, the combination of phases from different grid cell modules is adopted. If the spatial periods of all grid cell modules are pairwise co-prime, the total coding range $\Lambda$ is the product of all periods, i.e., $\Lambda = \prod_i^M \lambda^i = \bar{\lambda}^M$, where $M$ is the number of modules and $\lambda^i$ represents the spatial period of the $i$-th module [1]. Assuming each module contains $N_0$ grid cells, the coding efficiency of Phase Coding is calculated to be,

$$R_g = \frac{\ln \prod_{i=1}^M \lambda^i}{M N_0} = \frac{M \ln \bar{\lambda}}{M N_0} = \frac{\ln \bar{\lambda}}{N_0}. \tag{6}$$

We see that the efficiency of Phase Coding is independent of the number of grid cell modules $M$. As the number of grid cell modules (and hence the number of neurons) increases, the efficiency of Phase Coding remains to be a constant, indicating that Phase Coding is much more efficient than Localized Coding in terms of utilizing the encoding resource of neurons.

**Non-local errors of Phase Coding.** The vast coding range of grid cells, which grows exponentially with the number of modules, comes with the expense of high sensitivity to noise. As illustration, we consider Phase Coding has two grid cell modules. The phase combinations $(\phi^1, \phi^2)$ of two modules span a two-dimensional toroidal space with periodic boundaries (Figure S1c). A straight line in the position space is mapped into parallel lines in the phase plane according to the conversion rule given by Eq. (2). Notably, due to the periodicity of the phase space, two points that are far away in the position space can be mapped into two points that are close in the phase plane (comparing blue and red dots in Figure S1a,c). This indicates that a small perturbation in the phase space can incur a large shift in the position space, leading to a large non-local error (Figure S1d) [1].

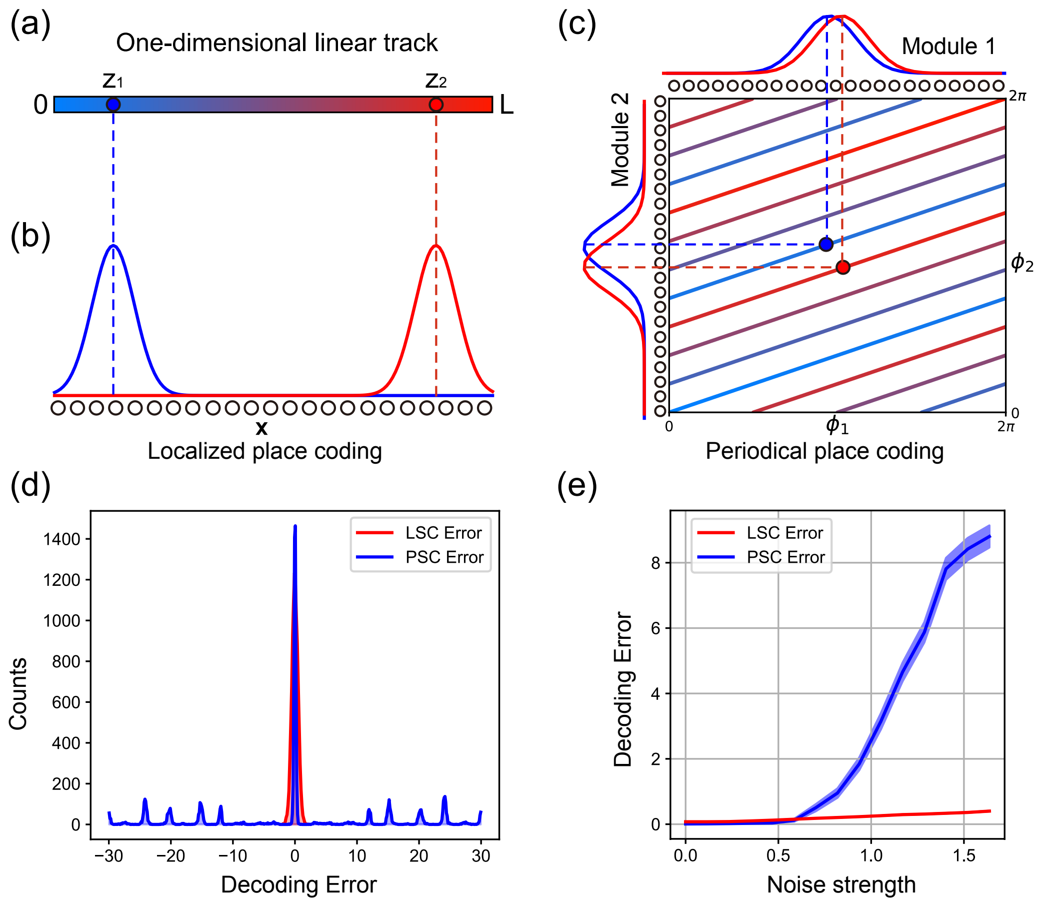


Figure S1: Illustrating the accuracy of Localized Coding and non-local errors of Phase Coding. **(a)**. A straight line representing the 1D linear track used in animal experiments, with colors representing continuously varying positions. The blue and red dots are two distant locations, denoted as $z_1$ and $z_2$, respectively. **(b)**. In Localized Coding, place cells are aligned according to their preferred positions, and the neuronal population activity forms a localized bump to encode a spatial location. The blue and red bumps encode the positions, $z_1$ and $z_2$, respectively. **(c)**. In Phase Coding, the phases of two grid cell modules ($\phi^1$, $\phi^2$) are combined to encode spatial locations. The straight line in (a) is mapped into parallel lines in the two-dimensional phase space. Note that two distant positions $z_1$ and $z_2$ in (a) are now mapped into two points (blue and red dots) close in the phase plane.

### 2 Decoding the animal position

We introduce the details of the methods used for decoding the animal location in this work.

#### 2.1 Gradient-based optimization of the posterior (GOP)

We have formulated the information integration between place and grid cells as a probabilistic inference problem. The posterior of $z$ and $\boldsymbol{\phi}$ given $\mathbf{r}_g$ and $\mathbf{r}_p$ is expressed as,

$$\begin{aligned} p(z, \boldsymbol{\phi}|\mathbf{r}_g, \mathbf{r}_p) &\propto p(\mathbf{r}_g|\boldsymbol{\phi})p(\mathbf{r}_p|z)p(\boldsymbol{\phi}, z) \\ &= \prod_{i=1}^{M} \left\{ p(\phi^i, z) \prod_{j=1}^{N_0} p\left[r_g(\theta_j^i)|\phi^i\right] \right\} \prod_{j=1}^{N} p\left[r_p(x_j)|z\right]. \end{aligned} \tag{7}$$

Maximizing the posterior is equivalent to maximizing the log posterior, and the latter is given by,

$$\begin{aligned}\ln p(z, \boldsymbol{\phi}|\mathbf{r}_g, \mathbf{r}_p) &\propto \ln p(\mathbf{r}_g|\boldsymbol{\phi}) + \ln p(\mathbf{r}_p|z) + \ln p(\boldsymbol{\phi}, z)\\ &= -\frac{\rho_g}{\sigma_{gi}^2}\sum_i \int_0^{2\pi} \left[r_g(\theta^i) - f_g(\theta^i;\phi^i)\right]^2 d\theta^i \\ &- \frac{\rho_p}{\sigma_p^2}\int_{-\infty}^{\infty}[r_p(x) - f_p(x;z)]^2 dx - \sum_i \frac{1}{\sigma_{\phi^i}^2}||\phi^i - \psi^i(z)||^2 + C.\end{aligned} \tag{8}$$

For the clarity of description, we approximate the discrete neuron distribution as a continuous one, i.e., $\sum_j [r(s_j) - f(s_j)]^2 = \rho_s \int [r(s) - f(s)]^2 ds$. $\rho_g = N_0/2\pi$ represents the density of grid cells and $\rho_p = N_p/L$ the density of place cells. $C = M \ln \sqrt{2\pi}\sigma_{gi} + M \ln \sqrt{2\pi}\sigma_{\phi^i} + N_p \ln \sqrt{2\pi}\sigma_p$ is a constant.

We calculate the gradients of each probability function with respect to $z$ and $\phi^i$. For $p(\boldsymbol{\phi}|z)$, the gradient of $\phi^i$ is calculated to be,

$$\begin{aligned}\frac{\partial}{\partial \phi^i}\ln p(\boldsymbol{\phi}, z) &= \frac{\partial}{\partial \phi^i}\left[-\frac{||\phi^i - \psi^i(z)||^2}{2\sigma_{\phi^i}^2}\right],\\ &= \frac{1}{\sigma_{\phi^i}^2}||\psi^i(z) - \phi^i||,\end{aligned} \tag{9}$$

and the gradient of $z$ is,

$$\begin{aligned}\frac{\partial}{\partial z}\ln p(\boldsymbol{\phi}, z) &= \frac{\partial}{\partial z}\sum_i\left[-\frac{||\phi^i - \psi^i(z)||^2}{2\sigma_{\phi^i}^2}\right],\\ &= \sum_i \frac{1}{\sigma_{\phi^i}^2}||\phi^i - \psi^i(z)||\frac{\partial \psi^i(z)}{\partial z},\\ &= \sum_i \frac{2\pi}{\lambda_i \sigma_{\phi^i}^2}||\phi^i - \psi^i(z)||.\end{aligned} \tag{10}$$

In the above, we have used the von-Mises distribution approximation of $p(\boldsymbol{\phi}, z)$ (see more details below).

For $p(\mathbf{r}_g|\boldsymbol{\phi})$, the gradient of $\phi^i$ is calculated to be,

$$\begin{aligned}\frac{\partial}{\partial \phi^i}\ln p(\mathbf{r}_g|\boldsymbol{\phi}) &= \frac{\partial}{\partial \phi^i}\int \rho_g\left[-\frac{(r_g(\theta^i)) - f_g(\theta^i;\phi^i))^2}{2\sigma_{gi}(\theta^i)}\right]d\theta^i,\\ &= \frac{\rho_g}{\sigma_{gi}^2}\int r_g(\theta^i)\frac{\partial f_g(\theta^i;\phi^i)}{\partial \phi^i}d\theta^i.\end{aligned} \tag{11}$$

The gradient of $z$ is calculated to be,

$$\frac{\partial}{\partial z}\ln p(\mathbf{r}_g|\boldsymbol{\phi}) = 0. \tag{12}$$

For $p(\mathbf{r}_p|z)$, the gradient of $\phi^i$ is calculated to be,

$$\frac{\partial}{\partial \phi^i}\ln p(\mathbf{r}_p|z) = 0, \tag{13}$$

and the gradient of $z$ is,

$$\begin{aligned}\frac{\partial}{\partial z}\ln p(\mathbf{r}_p|z) &= \frac{\partial}{\partial z}\rho_p\int\left[-\frac{(I_p(x) - f_p(x;z))^2}{2\sigma_p^2}\right]dx,\\ &= \frac{\rho_p}{\sigma_p^2}\int r_p(x)\frac{\partial f_p(x;z)}{\partial z}dx.\end{aligned} \tag{14}$$

Combining the above results, the gradients of the log posterior are obtained, which are,

$$\begin{aligned}\frac{\partial}{\partial z}\ln p(\boldsymbol{\phi}, z|\mathbf{r}_g, \mathbf{r}_p) &= \frac{\partial}{\partial z}\left[\ln p(\mathbf{r}_g|\boldsymbol{\phi}) + \ln p(\boldsymbol{\phi}, z) + \ln p(\mathbf{r}_p|z)\right],\\ &= \sum_i \frac{2\pi}{\lambda_i\sigma_{\phi^i}^2}||\phi^i - \psi^i(z)|| + \frac{\rho_p}{\sigma_p^2}\int r_p(x)\frac{\partial f_p(x;z)}{\partial z}dx,\end{aligned} \tag{15}$$

$$\frac{\partial}{\partial \phi^i} \ln p(\boldsymbol{\phi}, z|\mathbf{r}_g, \mathbf{r}_p) = \frac{\partial}{\partial \phi^i} \left[ \ln p(\mathbf{r}_g|\boldsymbol{\phi}) + \ln p(\boldsymbol{\phi}, z) + \ln p(\mathbf{r}_p|z) \right],$$
$$= \frac{1}{\sigma^2_{\phi^i}} ||\psi^i(z) - \phi^i|| + \frac{\rho_g}{\sigma^2_{gi}} \int r_g(\theta^i) \frac{\partial f_g(\theta^i; \phi^i)}{\partial \phi^i} d\theta^i. \tag{16}$$

The algorithm of gradient-based optimization of the posterior (GOP) is summarized in Table S1.

Table S1: The pseudo-code of GOP

| Step | Procedure |
|---|---|
| 1 | **Initialization:** Set initial values based on the historical information<br>$z^{(0)} = \hat{z}$, where $\hat{z}$ represents the decoded location at the previous time step.<br>$\boldsymbol{\phi}^{(0)} = \psi(z^{(0)})$ |
| 2 | **Gradient Ascent Update:** Iteratively update parameters using the gradients of the log posterior.<br>Update $z$: $z^{(k+1)} = z^{(k)} + \eta_z \nabla_z \ln p(z^{(k)}, \boldsymbol{\phi}^{(k)}|\mathbf{r}_g, \mathbf{r}_p)$<br>Update $\boldsymbol{\phi}$: $\boldsymbol{\phi}^{(k+1)} = \boldsymbol{\phi}^{(k)} + \eta_{\boldsymbol{\phi}} \nabla_{\boldsymbol{\phi}} \ln p(z^{(k)}, \boldsymbol{\phi}^{(k)}|\mathbf{r}_g, \mathbf{r}_p)$<br>where $\nabla_z \ln p$ and $\nabla_{\boldsymbol{\phi}} \ln p$ are the gradients of the log posterior with respect to $z$ and $\boldsymbol{\phi}$, and $\eta_z$ and $\eta_{\boldsymbol{\phi}}$ are the learning rates. |
| 3 | **Termination:** Stop iteration when a the local maximum is reached: $\nabla_z \ln p(z^{(k)}, \boldsymbol{\phi}^{(k)}|\mathbf{r}_g, \mathbf{r}_p) \approx 0$ and $\nabla_{\boldsymbol{\phi}} \ln p(z^{(k)}, \boldsymbol{\phi}^{(k)}|\mathbf{r}_g, \mathbf{r}_p) \approx 0$.<br>**Output:** $z, \boldsymbol{\phi}$ |

#### The von-Mises distribution approximation

In the main text, for convenience, we consider that $p(\boldsymbol{\phi}, z)$ satisfy Gaussian distributions. In reality, von-Mises distributions are more suitable to describe periodic variables. When computing gradients, we use von-Mises distributions to approximate Gaussian distributions, as the latter are not differentiable at the boundary.

Using the von-Mises distribution, the probability $p(\boldsymbol{\phi}, z)$ is written as,

$$p(\boldsymbol{\phi}, z) = \prod_{i=1}^{M} \frac{1}{2\pi I_0(\kappa)} \exp\left[\kappa \cos(\phi^i - \psi^i(z))\right]. \tag{17}$$

The modified Bessel function of the first kind and zero order $I_o(\kappa)$ serves as the normalizing constant, satisfying $\int_{-\pi}^{\pi} \exp\left[\kappa \cos(x - \mu)\right] = 2\pi I_o(\kappa)$. Parameters $\kappa$ and $\psi(z)$ are analogous to the variance and mean in the Gaussian distribution, respectively.

The logarithm of $p(\boldsymbol{\phi}, z)$ is expressed as,

$$\ln p(\boldsymbol{\phi}, z) = \sum_{i=1}^{M} \kappa \cos(\phi^i - \psi^i(z)) - M \ln(2\pi I_0(\kappa)), \tag{18}$$

where $\psi^i(z) = \text{mod}(z/\lambda^i) \times 2\pi = (z/\lambda^i) \times 2\pi - 2n^i\pi$, with $n^i = z//\lambda^i$ and the symbol $//$ represents the mathematical operation of divisibility. Substituting $\psi^i(z)$ into the above equation, we get,

$$\ln p(\boldsymbol{\phi}, z) = \sum_{i=1}^{M} \kappa \cos(\phi^i - \frac{z}{\lambda^i} 2\pi - 2n^i\pi) + C = \sum_{i=1}^{M} \kappa \cos(\phi^i - \frac{z}{\lambda^i} 2\pi) + C. \tag{19}$$

Thus, the gradient of $z$ is calculated to be,

$$\begin{aligned}
\frac{\partial}{\partial z}\ln p(\boldsymbol{\phi}, z) &= \frac{\partial}{\partial z}\sum_{i=1}^{M}\kappa\cos(\phi^i - \frac{z}{\lambda^i}2\pi), \\
&= \sum_i \frac{2\pi\kappa}{\lambda_i}\sin(\phi^i - \frac{z}{\lambda^i}2\pi), \\
&= \sum_i \frac{2\pi\kappa}{\lambda_i}\sin\left[\phi^i - \left(\frac{z}{\lambda^i}2\pi + 2n^i\pi\right)\right], \\
&= \sum_i \frac{2\pi\kappa}{\lambda_i}\sin(\phi^i - \psi^i(z)).
\end{aligned} \tag{20}$$

Given that the disparity between $\phi^i$ and $\psi^i(z)$ is sufficiently small, we can use the approximation $\sin(x) \approx x$ with $x$ sufficiently small. We have,

$$\frac{\partial}{\partial z}\ln p(\boldsymbol{\phi}, z) = \sum_i \frac{2\pi\kappa}{\lambda_i}||\phi^i - \psi^i(z)||. \tag{21}$$

Similarly, the gradient of $\phi^i$ is calculated to be

$$\frac{\partial}{\partial \phi^i}\ln p(\boldsymbol{\phi}, z) = \kappa||\psi^i(z) - \phi^i||. \tag{22}$$

These give the results in Eqs. (9,10).

#### 2.2 MAP based on grid cells' activity

Assuming independent Gaussian noises, the likelihood function of Phase Coding consists of two parts, which are:

$$p(\boldsymbol{\phi}, z) = \prod_{i=1}^{M} p(\phi^i, z) = \frac{1}{\sqrt{2\pi}\sigma_{\phi^i}}\exp\left[\sum_{i=1}^{M} -\frac{||\phi^i - \psi^i(z)||^2}{2\sigma_{\phi^i}^2}\right], \tag{23}$$

$$p(\mathbf{r}_g|\boldsymbol{\phi}) = \prod_{i=1}^{M}\prod_{j=1}^{N_g} p\left[r_g(\theta_j^i)|\phi^i\right] = \frac{1}{\sqrt{2\pi}\sigma_{gi}}\exp\left\{\sum_{i=1}^{M}\sum_{j=1}^{N_0} -\frac{\left[r_g(\theta_j^i) - f_g(\theta_j^i;\phi^i)\right]^2}{2\sigma_{gi}^2}\right\}, \tag{24}$$

where $\theta_j^i$ represents the preferred phase of the $j$th neuron in the $i$th module.

We can use maximum a posterior (MAP) to decode the position, which is written as,

$$\hat{z}, \hat{\boldsymbol{\phi}} = \arg\max_{z,\boldsymbol{\phi}} p(z, \boldsymbol{\phi}|\mathbf{r}_g) = \arg\max_{z,\boldsymbol{\phi}}\left[p(\mathbf{r}_g|\boldsymbol{\phi})p(\boldsymbol{\phi}, z)\right]. \tag{25}$$

### 3 Theoretical analysis of the model dynamics

#### 3.1 The model structure

We first introduce the details of the coupled network model.

The dynamics of place cells is given by,

$$\tau_p\frac{dU_p(x,t)}{dt} = -U_p(x,t) + \rho_p\int_0^L W_p(x,x')r_p(x',t)\,dx' + \sum_{i=1}^{M}\rho_g\int_{-\pi}^{\pi} W_{gi,p}(x,\theta^i)r_g(\theta^i,t)\,d\theta^i + I_p(x;z_0). \tag{26}$$

The dynamics of grid cells is given by,

$$\tau_g\frac{dU_g(\theta^i,t)}{dt} = -U_g(\theta^i,t) + \rho_g\int_{-\pi}^{\pi} W_g(\theta^i,\theta'^i)r_g(\theta'^i,t)\,d\theta'^i + \rho_p\int_0^L W_{gi,p}(x,\theta^i)r_p(x,t)\,dx + I_g(\theta^i;\phi_0^i). \tag{27}$$

The firing rate of neurons is given by,

$$r_s(s,t) = \frac{U_s(s,t)^2}{1 + k_s \rho_s \int U_s(s,t)^2 ds}, \quad s = x, \theta. \tag{28}$$

The recurrent connections between neurons in the P-CANN or G-CANN are given by

$$W_s(s,s') \quad = \quad \frac{J_s}{\sqrt{2\pi}a_s} \exp\left[ -\frac{||s-s'||^2}{2a_s^2} \right], \quad s = x, \theta. \tag{29}$$

The reciprocal connections between place and grid cells are given by,

$$W_{g,p}(x,\theta) = \frac{J_{g,p}}{\sqrt{2\pi}a_{gi,p}} \exp\left[ -\frac{||\theta - \psi(x)||^2}{2a_{gi,p}^2} \right], \tag{30}$$

where $\psi(x) = \mathrm{mod}(x/\lambda, 1) \times 2\pi - \pi$.

#### 3.2 The stationary states (bumps) of the coupled network

When the inhibition strength $k$ lies within a certain region [13], a CANN can hold a continuous family of bump-shape stationary states. For place cells, the bump state is expressed as,

$$\bar{U}_p(x) = A_p \exp\left[ -\frac{(x-z)^2}{4a_p^2} \right], \quad \bar{r}_p(x) = \hat{r}_p \exp\left[ -\frac{(x-z)^2}{2a_p^2} \right]. \tag{31}$$

For grid cells, the bump state is expressed as,

$$\bar{U}_g(\theta^i) = A_g \exp\left[ -\frac{||\theta^i - \phi^i||^2}{4a_p^2} \right], \quad \bar{r}_g(\theta^i) = \hat{r}_g \exp\left[ -\frac{||\theta^i - \phi^i||^2}{2a_{gi}^2} \right]. \tag{32}$$

In the above $A_p$ and $A_g$ represent the bump heights in P-CANN and G-CANN, respectively, and $\hat{r}_p$ and $\hat{r}_g$ the maximum firing rates of place cells and grid cells, respectively. We check the condition when the coupled networks hold these bump states as their stationary states. For clearance, hereafter, we denote the Gaussian function as $\mathcal{N}(s|s', a) = \exp\left[ -||s - s'||^2/4a^2 \right]$.

Substituting Eqs. (31 ,32) into the grid cell dynamics, we get,

$$\begin{aligned} &\tau_g \left[ A_g \frac{||\theta^i - \phi^i||}{2a_{gi}^2} \frac{d\phi^i}{dt} + \tau_g \frac{dA_g}{dt} \right] \mathcal{N}(\theta|\phi^i, a_{gi}) = (-A_g + \frac{\rho_g J_g}{\sqrt{2}} \hat{r}_g) \mathcal{N}(\theta|\phi^i, a_{gi}) \\ &+ \rho_p \int \frac{J_{g,p}\hat{r}_p}{\sqrt{2\pi}a_{gi,p}} \mathcal{N}(\theta|\psi(x), \frac{a_{gi,p}}{\sqrt{2}}) \mathcal{N}(x|z, \frac{a_p}{\sqrt{2}})] dx + I_g^i(\theta^i, t), \end{aligned} \tag{33}$$

where $\mathcal{N}(\theta|\phi^i, a_{gi}) = \exp\left[ -||\theta^i - \phi^i||^2/4a_{gi}^2 \right]$. By setting $I_g^i = 0$ and $dA_g/dt = 0$, the dynamics of grid cells can be simplified as,

$$(A_g - \frac{\rho_g J_g}{\sqrt{2}} \hat{r}_g) \mathcal{N}(\theta|\phi^i, a_{gi}) = \rho_p \int \frac{J_{g,p}\hat{r}_p}{\sqrt{2\pi}a_{gi,p}} \mathcal{N}(\theta|\psi(x), \frac{a_{gi,p}}{\sqrt{2}}) \mathcal{N}(x|z, \frac{a_p}{\sqrt{2}}) dx. \tag{34}$$

To calculate the integral in the right-hand side of Eq. (34), we need to map the bump from the phase representation to the location representation. Because of the periodic nature, a phase corresponds to many locations. Denote $\psi^{-1}(\theta^i; z) = (\lambda^i/2\pi)\theta^i + n^i(z)$ to be the location corresponding to phase $\theta$ closest to $z$. We have,

$$\begin{aligned} \mathcal{N}(\theta^i|\psi^i(z), a_{gi}) &= \exp\left[ -\frac{||\theta^i - \psi^i(z)||^2}{2a_{gi}^2} \right], \\ &= \exp\left[ -\frac{||\theta^i - 2\pi(z/\lambda^i - n^i)||^2}{2a_{gi}^2} \right], \\ &= \exp\left[ -\frac{((\lambda^i/2\pi)\theta^i + n^i(z) - z)^2}{2a_p^2} \right], \\ &= \exp\left[ -\frac{(\psi^{-1}(\theta^i; z) - z)^2}{2a_p^2} \right], \\ &= \mathcal{N}(z|\psi^{-1}(\theta^i; z), a_p). \end{aligned} \tag{35}$$

Note that $|\psi^{-1}(\theta^i; z) - z|$ must be smaller than $\lambda^i/2$, implying that $||\theta^i - 2\pi(z/\lambda^i - n^i(z))|| = |\theta^i - 2\pi(z/\lambda^i - n^i(z))| < \pi$, so that the operation of circular distance can be removed from the above equation.

Using this property, we can further simplify Eq. (34) as,

$$
\begin{aligned}
(A_g - \frac{\rho_g J_g}{\sqrt{2}}\hat{r}_g)\mathcal{N}(\theta|\phi^i, a_{gi}) &= \rho_p \frac{J_{g,p}\hat{r}_p}{\sqrt{2\pi}a_{gi,p}} \int \mathcal{N}(\theta|\psi(x), \frac{a_{gi,p}}{\sqrt{2}})\mathcal{N}(x|z, \frac{a_p}{\sqrt{2}})dx, \\
&= \rho_p \frac{J_{g,p}\hat{r}_p}{\sqrt{2\pi}a_{gi,p}} \int \mathcal{N}(z|\psi^{-1}(\theta; x), \frac{\lambda}{2\pi}\frac{a_{gi,p}}{\sqrt{2}})\mathcal{N}(x|z, \frac{a_p}{\sqrt{2}})dx.
\end{aligned}
\tag{36}
$$

We see that, the condition for the above equation to be satisfied is,

$$
a_{gi,p} = a_{gi} = a_p(2\pi/\lambda), \tag{37}
$$

We substitute this condition into the above equation and get,

$$
\begin{aligned}
(A_g - \frac{\rho_g J_g}{\sqrt{2}}\hat{r}_g)\mathcal{N}(\theta|\phi^i, a_{gi}) &= \frac{\rho_p J_{g,p}\hat{r}_p}{\sqrt{2}}\mathcal{N}(x|\psi^{-1}(\theta; x), a_p), \\
&= \frac{\rho_p J_{g,p}\hat{r}_p \lambda}{2\sqrt{2}\pi}\mathcal{N}(\theta|\psi(z), a_{gi}).
\end{aligned}
\tag{38}
$$

Therefore, in the steady state, the bump center of grid cell modules is $\phi^i = \psi(z)$, with $z$ the bump center of place cells.

Substituting Eqs. (31 ,32) into the place cell dynamics, we get,

$$
\begin{aligned}
\tau_p A_p \frac{x-z}{2a_p^2}\mathcal{N}(z, 2a_p^2)\frac{dz}{dt} + \tau_p \frac{dA_p}{dt}\mathcal{N}(z, 2a_p^2) =& (-A_p + \frac{\rho_p J_p}{\sqrt{2}}\hat{r}_p)\mathcal{N}(z, 2a_p^2) \\
&+ \sum_{i=1}^{M} \frac{\rho_g J_{g,p}\hat{r}_g}{\sqrt{2}} \exp\left[-\frac{||\psi(x) - \phi^i||^2}{4a_{gi}^2}\right] + I_{\mathrm{p}}(x, t).
\end{aligned}
\tag{39}
$$

The divisive normalization can be simplified as,

$$
\hat{r}_s = \frac{A_s^2}{1 + \sqrt{2\pi}a_s\rho_s k_s A_s^2}, \tag{40}
$$

where $s$ denotes the cell type.

We calculate the stationary state of the networks when no external input exist ($\mathbf{I}_p = \mathbf{I}_g^i = 0$). By setting $dA_p/dt = dA_g/dt = 0$ and $dz/dt = d\phi^i/dt = 0$, we obtain

$$
A_p = \frac{\rho_p J_p}{\sqrt{2}}\hat{r}_p + \frac{1}{\sqrt{2}}\sum_i \rho_g \hat{r}_g J_{g,p}, \tag{41}
$$

$$
A_g = \frac{\rho_g J_g}{\sqrt{2}}\hat{r}_g + \frac{1}{\sqrt{2}}\rho_p r_p J_{g,p}. \tag{42}
$$

Using divisive normalization, we get,

$$
A_g = \frac{\rho_g J_g}{\sqrt{2}} \frac{A_g^2}{1 + \sqrt{2\pi}a_{gi}\rho_g k_g A_g^2} + \frac{\rho_p J_{g,p}}{\sqrt{2}} \frac{A_p^2}{1 + \sqrt{2\pi}a_p\rho_p k_p A_p^2}, \tag{43}
$$

$$
A_p = \frac{\rho_p J_p}{\sqrt{2}} \frac{A_p^2}{1 + \sqrt{2\pi}a_p\rho_p k_p A_p^2} + \frac{1}{\sqrt{2}}\sum_i \frac{\rho_g J_{g,p} a_p}{a_{gi}} \frac{A_g^2}{1 + \sqrt{2\pi}a_{gi}\rho_g k_g A_g^2}. \tag{44}
$$

Previous research has shown that for CANNs, external inputs primarily influence the network dynamics by altering the position of the network's activity bump [14; 13]. This sensitivity arises because a CANN exhibits neutral stability in the direction of the activity bump's movement, making them highly responsive to external inputs. In contrast, the height of the activity bump is less sensitive to changes in external inputs. When examining changes in bump height, we consider that recurrent inputs within each network are much stronger than reciprocal ones, allowing us to disregard reciprocal inputs and simplify the above expressions,

$$A_g = \frac{\rho_g J_g}{\sqrt{2}} \frac{A_g^2}{1+\sqrt{2\pi} a_{gi} \rho_g k_g A_g^2}, \tag{45}$$

$$A_p = \frac{\rho_p J_p}{\sqrt{2}} \frac{A_p^2}{1+\sqrt{2\pi} a_p \rho_p k_p A_p^2}. \tag{46}$$

We have,

$$A_g = \frac{1}{4\sqrt{\pi} a_{gi} \rho_g k_g} \left( \rho_g J_g + \sqrt{\rho_g^2 J_g^2 - 8\sqrt{2\pi} a_{gi} \rho_g k_g} \right), \tag{47}$$

$$A_p = \frac{1}{4\sqrt{\pi} a_p \rho_p k_p} \left( \rho_p J_p + \sqrt{\rho_p^2 J_p^2 - 8\sqrt{2\pi} a_p \rho_p k_p} \right). \tag{48}$$

#### 3.3 Simplifying the model dynamics by the projection method

In a CANN, its stationary states constitute a neutrally stable sub-manifold, implying that the network dynamics can be effectively represented by a small number of dominant motion modes, such as the variations in height and position of the stationary states. Therefore, by projecting the network dynamics onto these dominant motion modes, we can simplify the network dynamics significantly. The first two dominating motion modes of a CANN can be expressed as,

$$\text{height} : u_s^0(s) = \bar{U}_s(s), \tag{49}$$

$$\text{position} : u_s^1(s) = \frac{\partial \bar{U}_s(s)}{\partial s}. \tag{50}$$

##### 3.3.1 The simplified dynamics of place cells

For place cells, the first two modes are written as,

$$u_p^0(x) = \exp\left[ -\frac{(x-z)^2}{4a_p^2} \right], \tag{51}$$

$$u_p^1(x) = [x-z] \exp\left[ -\frac{(x-z)^2}{4a_p^2} \right]. \tag{52}$$

We consider that the field width of place cells is much smaller than the space range, i.e., $a_p \ll L$, and the integral from $-L/2$ to $L/2$ can be approximate as from $-\infty$ to $\infty$.

For the projection on $u_p^0$, the result is,

$$\begin{aligned} \text{Left} &= \int \tau_p \frac{A_p}{2a_p^2} \frac{dz}{dt} (x-z) \exp\left[ -\frac{(x-z)^2}{4a_p^2} \right] u_p^0(x) dx + \int \tau_p \frac{dA_p}{dt} \exp\left[ -\frac{(x-z)^2}{4a_p^2} \right] u_p^0(x) dx, \\ &= \sqrt{2\pi} a_p \tau_p \frac{dA_p}{dt}. \end{aligned} \tag{53}$$

$$\begin{aligned} \text{Right} &= \int \left( -A_p + \frac{\rho_p J_p}{\sqrt{2}} \hat{r}_p \right) \exp\left[ -\frac{(x-z)^2}{4a_p^2} \right] u_p^0(x) dx \\ &+ \int \sum_i \frac{\rho_g \hat{r}_g J_{g,p}}{\sqrt{2}} \exp\left[ -\frac{||\phi^i - \psi^i(x)||^2}{4a_{gi}^2} \right] u_p^0(x) dx + \int I_{\text{p}}(x,t) u_p^0(x) dx, \\ &= \sqrt{2\pi} a_p (-A_p + \frac{\rho_p J_p}{\sqrt{2}} \hat{r}_p) + \sqrt{\pi} a_p \sum_i \rho_g \hat{r}_g J_{g,p} \exp\left[ -\frac{||\phi^i - \psi^i(z)||^2}{8a_{gi}^2} \right] + \int I_p u_p^0 dx. \end{aligned} \tag{54}$$

For the projection on $u_p^1$, the result is,

$$\begin{aligned}\text{Left} &= \int \tau_p A_p \frac{x-z}{2a_p^2}\frac{dz}{dt}\exp\left[-\frac{(x-z)^2}{4a_p^2}\right]u_p^1(x)dx \\ &+ \int \tau_p \frac{dA_p}{dt}\exp\left[-\frac{(x-z)^2}{4a_p^2}\right]u_p^1(x)dx, \\ &= 2\sqrt{\pi}a_p\tau_p A_p\frac{dz}{dt}.\end{aligned} \tag{55}$$

$$\begin{aligned}\text{Right} &= \int \left(-A_p + \frac{\rho_p J_p}{\sqrt{2}}\hat{r}_p\right)\exp\left[-\frac{(x-z)^2}{4a_p^2}\right]u_p^1(x)dx \\ &+ \int \sum_i \frac{\rho_g \hat{r}_g J_{g,p}}{\sqrt{2}}\exp\left[-\frac{[x-\psi^{-1}(\phi^i)]^2}{4a_p^2}\right]u_p^1(x)dx + \int I_{\mathrm{p}}(x,t)u_p^1(x)dx \\ &= \frac{\sqrt{\pi}a_p}{2}\sum_i \frac{\lambda_i J_{g,p}\rho_g\hat{r}_g}{2\pi}||\phi^i-\psi^i(z)||\exp\left[-\frac{||\psi^i(z)-\phi^i||^2}{8a_{gi}^2}\right] + \int I_p u_p^1 dx\end{aligned} \tag{56}$$

Combining the above equalities, we obtain

$$\frac{dA_p}{dt} = \frac{1}{\tau_p}\left\{-A_p + \frac{\rho_p J_p}{\sqrt{2}}\hat{r}_p + \frac{1}{\sqrt{2}}\sum_i \rho_g\hat{r}_g J_{g,p}\exp\left[-\frac{||\psi^i(z)-\phi^i||^2}{8a_{gi}^2}\right] + \frac{1}{\sqrt{2\pi}a_p}\int I_p u_p^0 dx\right\}, \tag{57}$$

$$\frac{dz}{dt} = \frac{1}{\tau_p A_p}\left\{\frac{1}{4}\sum_i \frac{\lambda_i}{2\pi}J_{g,p}\rho_g\hat{r}_g||\phi^i-\psi^i(z)||\exp\left[-\frac{||\psi^i(z)-\phi^i||^2}{8a_{gi}^2}\right] + \frac{1}{2\sqrt{\pi}a_p}\int I_p u_p^1 dx\right\}. \tag{58}$$

##### 3.3.2 The simplified dynamics of grid cells

For grid cells, the first two dominant motion modes are written as,

$$u_g^0(\theta^i) = \exp\left[-\frac{||\theta^i-\phi^i||^2}{4a_{gi}^2}\right], \tag{59}$$

$$u_g^1(\theta^i) = ||\theta^i-\phi^i||\exp\left[-\frac{||\theta^i-\phi^i||^2}{4a_{gi}^2}\right]. \tag{60}$$

For the projection on $u_g^0$, the result is,

$$\begin{aligned}\text{Left} &= \int \left\{\tau_g A_g \frac{||\theta^i-\phi^i||}{2a_{gi}^2}\exp\left[-\frac{||\theta^i-\phi^i||^2}{4a_{gi}^2}\right]\frac{d\phi^i}{dt}\right\}u_g^0(\theta^i)d\theta^i \\ &+ \int \left\{\tau_g\frac{dA_g}{dt}\exp\left[-\frac{||\theta^i-\phi^i||^2}{4a_{gi}^2}\right]\right\}u_g^0(\theta^i)d\theta^i, \\ &= \sqrt{2\pi}a_{gi}\tau_g\frac{dA_g}{dt}.\end{aligned} \tag{61}$$

$$\begin{aligned}\text{Right} &= \int \left(-A_g + \frac{\rho_g J_g}{\sqrt{2}}\right)\hat{r}_g\exp\left[-\frac{||\theta^i-\phi^i||^2}{4a_{gi}^2}\right]u_g^0(\theta^i)d\theta^i \\ &+ \int \frac{\rho_p\hat{r}_p J_{g,p}}{\sqrt{2}}\frac{\lambda_i}{2\pi}\exp\left[-\frac{||\theta^i-\psi^i(z)||^2}{4a_{gi}^2}\right]u_g^0(\theta^i)d\theta^i + \int I_{\mathrm{g}}(\theta^i,t)u_g^0(\theta^i)d\theta^i, \\ &= \sqrt{2\pi}a_{gi}\left(-A_g + \frac{\rho_g J_g}{\sqrt{2}}\hat{r}_g\right) + \sqrt{\pi}a_p\rho_p\hat{r}_p J_{g,p}\exp\left[-\frac{||\phi^i-\psi^i(z)||^2}{8a_{gi}^2}\right] + \int I_g^i u_g^0 dx.\end{aligned} \tag{62}$$

For the projection on $u_g^1$, the result is,

$$\begin{aligned}
\text{Left} &= \int \left\{ \tau_g A_g \frac{||\theta^i - \phi^i||}{2a_{gi}^2} \exp\left[ -\frac{||\theta^i - \phi^i||^2}{4a_{gi}^2} \right] \frac{d\phi^i}{dt} \right\} u_g^1(\theta^i) d\theta^i \\
&+ \int \left\{ \tau_g \frac{dA_g}{dt} u_g^0(\theta^i) \right\} u_g^1(\theta^i) d\theta^i, \\
&= 2\sqrt{\pi} a_{gi} \tau_g A_g \frac{d\phi^i}{dt}.
\end{aligned} \tag{63}$$

$$\begin{aligned}
\text{Right} &= \int \left( -A_g + \frac{\rho_g J_g}{\sqrt{2}} \hat{r}_g \right) \exp\left[ -\frac{||\theta^i - \phi^i||^2}{4a_{gi}^2} \right] u_g^1(\theta^i) d\theta^i \\
&+ \int \frac{a_p}{a_{gi}} \frac{\rho_p \hat{r}_p J_{g,p}}{\sqrt{2}} \exp\left[ -\frac{||\theta^i - \psi^i(z)||^2}{4a_{gi}^2} \right] u_g^1(\theta^i) d\theta^i \\
&+ \int I_{\mathrm{g}}(\theta^i, t) u_g^1(\theta^i) d\theta^i, \\
&= \frac{\sqrt{\pi} a_p}{2} J_{g,p} \rho_p \hat{r}_p ||\phi^i - \psi^i(z)|| \exp\left[ -\frac{||\psi^i(z) - \phi^i||^2}{8a_{gi}^2} \right] + \int I_g^i u_g^1 dx.
\end{aligned} \tag{64}$$

Synthesizing the above equalities, we obtain

$$\frac{dA_g}{dt} = \frac{1}{\tau_g} \left\{ -A_g + \frac{\rho_g J_g}{\sqrt{2}} \hat{r}_g + \frac{1}{\sqrt{2}} \rho_p \hat{r}_p J_{g,p} \exp\left[ -\frac{||\phi^i(t) - \psi^i(z)||^2}{8a_{gi}^2} \right] + \frac{1}{\sqrt{2\pi} a_{gi}} \int I_g^i u_g^0 dx \right\}. \tag{65}$$

$$\frac{d\phi^i}{dt} = \frac{1}{\tau_g A_g} \left\{ \frac{\lambda}{8\pi} J_{g,p} \rho_p \hat{r}_p ||\psi^i(z) - \phi^i|| \exp\left[ -\frac{||\phi^i(t) - \psi^i(z)||^2}{8a_{gi}^2} \right] + \frac{1}{2\sqrt{\pi} a_{gi}} \int I_g^i u_g^1 dx \right\}. \tag{66}$$

Combing the above results and assuming that the network bump centers are close enough, i.e., $||\phi^i(t) - \psi^i(z)||$ are sufficiently small, we get the final form of the simplified network dynamics, which are,

$$\begin{aligned}
\frac{dA_p}{dt} &= \frac{1}{\tau_p} \left\{ -A_p + \frac{\rho_p J_p}{\sqrt{2}} \hat{r}_p + \frac{1}{\sqrt{2}} \sum_i \rho_g \hat{r}_g J_{g,p} + \frac{1}{\sqrt{2\pi} a_p} \int I_p u_p^0 dx \right\}, && (67) \\
\frac{dz}{dt} &= \frac{1}{\tau_p A_p} \left\{ \frac{1}{4} \sum_i \frac{\lambda_i}{2\pi} J_{g,p} \rho_g \hat{r}_g ||\phi^i - \psi^i(z)|| + \frac{1}{2\sqrt{\pi} a_p} \int I_p u_p^1 dx \right\} && (68) \\
\frac{dA_g}{dt} &= \frac{1}{\tau_g} \left\{ -A_g + \frac{\rho_g J_g}{\sqrt{2}} \hat{r}_g + \frac{1}{\sqrt{2}} \rho_p \hat{r}_p J_{g,p} + \frac{1}{\sqrt{2\pi} A_g} \int I_g^i u_g^0 dx \right\}, && (69) \\
\frac{d\phi^i}{dt} &= \frac{1}{\tau_g A_g} \left\{ \frac{\lambda}{8\pi} J_{g,p} \rho_p \hat{r}_p ||\psi^i(z) - \phi^i|| + \frac{1}{2\sqrt{\pi} a_{gi}} \int I_g^i u_g^1 dx \right\}. && (70)
\end{aligned}$$

##### 3.4 The coupled network implementing GOP

From Eqs.(15,16), the gradients of the logarithm posterior are given by,

$$\frac{\partial}{\partial z} \ln p(\boldsymbol{\phi}, z | \mathbf{r}_g, \mathbf{r}_p) = \sum_i \frac{2\pi}{\lambda_i \sigma_{\phi^i}^2} ||\phi^i - \psi^i(z)|| + \frac{\rho_p}{\sigma_p^2} \int I_p(x) \frac{\partial f_p(x; z)}{\partial z} dx, \tag{71}$$

$$\frac{\partial}{\partial \phi^i} \ln p(\boldsymbol{\phi}, z | \mathbf{r}_g, \mathbf{r}_p) = \frac{1}{\sigma_{\phi^i}^2} ||\psi^i(z) - \phi^i|| + \frac{\rho_g}{\sigma_{gi}^2} \int r_g(\theta^i) \frac{\partial f_g(\theta^i; \phi^i)}{\partial \phi^i} d\theta^i. \tag{72}$$

From Eqs. (68,70), the dynamics of the bump centers of place and grid cells are written as,

$$\frac{dz}{dt} = \frac{1}{\tau_p A_p}\left\{\frac{1}{4}\sum_i \frac{\lambda_i}{2\pi} J_{g,p}\rho_g \hat{r}_g ||\phi^i - \psi^i(z)|| + \frac{1}{2\sqrt{\pi}a_p}\int I_p u_p^1 dx\right\}, \tag{73}$$

$$\frac{d\phi^i}{dt} = \frac{1}{\tau_g A_g}\left\{\frac{\lambda}{8\pi} J_{g,p}\rho_p \hat{r}_p ||\psi^i(z) - \phi^i|| + \frac{1}{2\sqrt{\pi}a_{gi}}\int I_g^i u_g^1 d\theta\right\}. \tag{74}$$

We consider the external inputs to the networks are given by, $\mathbf{I}_g^i = \alpha_{gi}\mathbf{r}_g$ and $\mathbf{I}_p = \alpha_p \mathbf{r}_p$, where $\alpha_{gi}$ and $\alpha_p$ represent the strengths of external inputs to the grid cells and place cells, respectively.

Comparing the gradients of the posterior Eqs. (71,72) with the network dynamics Eqs. (73,74), we see that they are equivalent if the below parameter conditions are satisfied, which are,

$$(\frac{\lambda_i}{2\pi})^2 \frac{J_{g,p}\rho_g \hat{r}_g}{4A_p\tau_p} = \frac{1}{\sigma_{\phi^i}^2}, \qquad \frac{a_p\alpha_p}{\sqrt{\pi}A_p^3\rho_p\tau_p} = \frac{1}{\sigma_p^2},$$
$$\frac{\lambda_i}{2\pi}\frac{J_{g,p}\rho_p\hat{r}_p}{4A_g\tau_g} = \frac{1}{\sigma_{\phi^i}^2}, \qquad \frac{a_{gi}\alpha_{gi}}{\sqrt{\pi}A_g^3\rho_g\tau_g} = \frac{1}{\sigma_{gi}^2}. \tag{75}$$

From the above equalities, we find the necessary condition for $\sigma_{\phi^i}$ well-defined is $2\pi A_p\tau_p\rho_p\hat{r}_p = \lambda_i A_g^i \tau_g \rho_g^i \hat{r}_g^i$. The relationships between the parameters in the probabilistic model and those in the network dynamics are: $\sigma_p^2 = (\sqrt{\pi}A_p^3\rho_p\tau_p)/(2a_p\alpha_p)$, $\sigma_{\phi^i}^2 = (8\pi A_g\tau_g)/(\lambda_i J_{g,p}\rho_p\hat{r}_p)$ and $\sigma_{gi}^2 = (\sqrt{\pi}A_g^3\rho_g\tau_g)/(2a_{gi}\alpha_{gi})$.

For the sake of simulation convenience, we can simplify the above parameter constraint by decomposing them into a more manageable form. Assuming the neuron densities and recurrent strength of grid cells and place cells are equal, i.e. $\rho_p = \rho_g$, $J_g = J_p$, and that the products of interaction width and global inhibition strength within each type of cells are the same, $a_{gi}k_g = a_p k_p$. Therefore, according to the previous derivation of height of stationary synaptic input $A_s$ and maximal firing rate $\hat{r}_s$, we have

$$A_s = \frac{\rho_s J_s}{\sqrt{2}}\frac{A_s^2}{1+\sqrt{2\pi}a_s\rho_s k_g A_s^2} + \frac{\rho_p J_{g,p}}{\sqrt{2}}\frac{A_s^2}{1+\sqrt{2\pi}a_s\rho_p k_p A_s^2}, \tag{76}$$

$$\hat{r}_s = \frac{A_s^2}{1+\sqrt{2\pi}a_s\rho_s k_s A_s^2}, \tag{77}$$

where s denotes the cell type. We see that $A_p = A_g$ and $\hat{r}_p = \hat{r}_g$. Substituting into well-defined condition of $\sigma_{\phi^i}$, $2\pi A_p\tau_p\rho_p\hat{r}_p = \lambda_i A_g^i\tau_g\rho_g^i\hat{r}_g^i$, we can conclude that $a_p\tau_p = a_{gi}\tau_g$. Utilizing the relation between interaction width $a_{gi}/2\pi = a_p/\lambda_i$, we can decompose the constrain as

$$A_p = A_g, \quad J_p = J_g, \quad \frac{k_p}{2\pi} = \frac{k_g}{\lambda_i}, \quad \tau_p = \tau_g\frac{\lambda_i}{2\pi}. \tag{78}$$

##### 3.5 The energy function of the model dynamics

The simplified model dynamics has a energy function, which is the negative posterior $E = -\ln P(z, \boldsymbol{\phi}|\mathbf{r}_p, \mathbf{r}_g)$, that is,

$$\frac{dE}{dt} = -\sum_i \frac{\partial E}{\partial y_i}\frac{dy_i}{dt} = -\sum_i (\frac{dy_i}{dt})^2 \le 0, \qquad y_1 = z, \quad y_2 = \boldsymbol{\phi}. \tag{79}$$

#### 4 Simulation experiments for evaluating network performances

We introduce the details of implementing the network decoding, GOP, and MAP to decode animal position given either both environmental and self-motion cues ($I_p > 0$, $I_g > 0$) or only the self-motion cue ($I_p = 0$, $I_g > 0$).

First, we consider a 1D spatial range of $L = 60$, with the animal positioned at $L/2$. For MAP, we directly compute the maximum of the posterior probability distribution given by Eq. (23). For both network decoding and GOP, they require an initial state that reflects the animal's estimated position from the previous time step. Since the animal moves continuously through the space, we set the initial position at $z_0 = L/2 - 0.5$, close to the actual animal position ($L/2$). We then run the network dynamics (as described in Eqs. (2–3)) or the GOP algorithm (as described in Eqs. (8–9)) for 2000 iterative steps. At each time step, for the network decoding, the animal's location is read out based on the center of place cells' activity, which is given by,

$$z(t) = \frac{\sum_i r_p(x_i, t) x_i}{\sum_i r_p(x_i, t)}, \tag{80}$$

where $r_p(x_i, t)$ represents the activity of place cells with preferred position at $x_i$ at time t.

For the probabilistic inference model (both MAP and GOP), calculating the posterior requires the widths of the tuning curves $a_p$ and $a_g$, the noise strengths $\sigma_p$ and $\sigma_g$, and the standard deviation $\sigma_\phi$ of the correlation prior between position and phase. Table S2 lists the default values for these parameters. For experiments that involve varying $\sigma_p$ and $\sigma_g$, the corresponding values are shown in the respective figures.

| Variable | Value | Description |
|---|---|---|
| $a_{g1}$ | 0.63 | Tuning curve width of grid cells (module 1) |
| $a_{g2}$ | 0.47 | Tuning curve width of grid cells (module 2) |
| $a_{g3}$ | 0.38 | Tuning curve width of grid cells (module 3) |
| $\sigma_{\phi^1}$ | 0.25 | Std. dev. of correlation prior (module 1) |
| $\sigma_{\phi^2}$ | 0.19 | Std. dev. of correlation prior (module 2) |
| $\sigma_{\phi^3}$ | 0.15 | Std. dev. of correlation prior (module 3) |
| $a_p$ | 0.3 | Tuning curve width of place cells |
| $\sigma_p$ | 0.25 | Noise strength of place cell inputs |
| $\sigma_g$ | 0.2 | Noise strength of grid cell inputs |

Table S2: Default parameters of the probabilistic inference model.

For the coupled network, the parameters include: the connection width of P-CANN ($a_p$), the connection width of each G-CANN ($a_{gi}$), the connection strength of P-CANN ($J_p$), the connection strength of all G-CANNs ($J_g$), the connection strength between P-CANN and G-CANNs ($J_{pg}$), the global inhibition strength of P-CANN ($k_p$), the global inhibition strength of each G-CANN ($k_{gi}$), the time constant of P-CANN ($\tau_p$), the time constant of each G-CANN ($\tau_{gi}$), the number of place cell neurons ($N_p$), the number of grid cell neurons within each module ($N_g$), the number of grid cell modules ($M$), and the spacings of each G-CANN ($\lambda_i$). The values of these parameters are shown in Table S3.

| Variable | Value | Variable | Value | Variable | Value | Variable | Value |
|---|---|---|---|---|---|---|---|
| $a_p$ | 0.3 | $a_{g1}$ | 0.63 | $a_{g2}$ | 0.47 | $a_{g3}$ | 0.38 |
| $k_p$ | 20 | $k_{g,1}$ | 9.55 | $k_{g,2}$ | 12.73 | $k_{g,3}$ | 15.91 |
| $N_p$ | 200 | $N_g$ | 20 | $J_p, J_g$ | 20 | $J_{g,p}$ | 0.8 |
| $M$ | 3 | $\lambda_1$ | 3 | $\lambda_2$ | 4 | $\lambda_3$ | 5 |
| $\tau_p$ | 1.0 | $\tau_{g,1}$ | 2.09 | $\tau_{g,2}$ | 1.57 | $\tau_{g,3}$ | 1.26 |

Table S3: Parameters of the coupled network encoding a single map.

#### 4.1 Position decoding over time

In the above, we described how to decode position at a single time step. To further validate the network's ability of robustly representing position through path integration, thereby enhancing grid cell robustness and eliminating non-local errors when only the self-motion cue is available ($I_p = 0$), we compare the decoding performance of the network and MAP when the animal moves continuously.

Without loss of generality, we assume that the animal moves at a constant speed in a linear track. The initial position is set as $z(t = 0) = 0$, and at each step the animal moves $\Delta z = 0.1$ for totally 20 steps. At each step, We read out the bump center of each grid module $\phi^i(t)$, and the bump center of P-CANN $z(t)$. $z(t)$ is regarded as the decoded position.

At each step, the animal does not directly perceive its absolute position $z$ but only the displacement $\Delta z$. Since grid cells perform path integration, at time $t+1$ the physical displacement $\Delta z$ is first mapped to the phase space of each grid module, yielding $\Delta\phi^i = \Delta z\,/\,(2\pi) \times \lambda_i$. The network input for time $t+1$ is then generated for each grid cell centered at $\phi^i(t+1) = \phi^i(t) + \Delta\phi^i$. Note that in our simulation of path integration with only the self-motion cue, decoding error in $\phi$ from previous steps accumulates over time.

We performed 1000 independent experiments, each producing a trajectory of decoded positions. At each time step, we calculated the deviation from the decoded position to the true animal position and then calculated the mean and variance of these errors.

##### 4.2 Replicating experimental findings on remapping phenomena of place cells

The simulation results for the inhibited grid cell condition are shown in Figure 6 in the main text, while the results for the stimulated grid cell condition are presented in Figure S2.

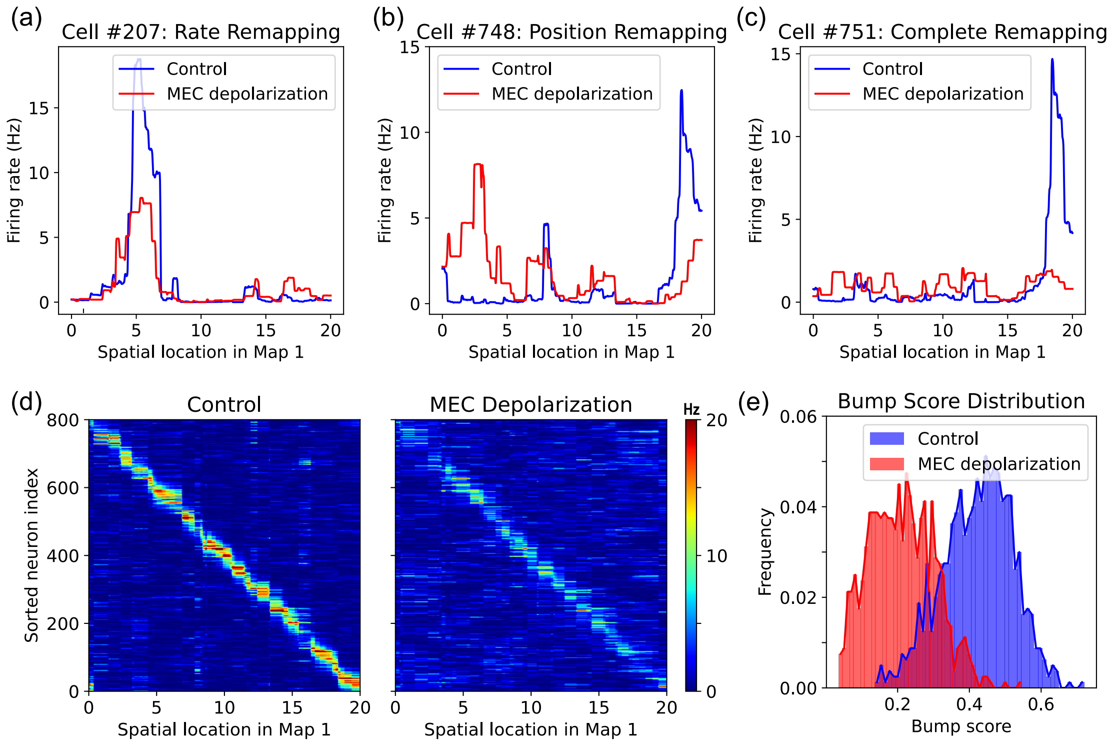


Figure S2: Depolarization of grid cell activity induces place cell remapping. **(a-c)**. Tuning curves of three place cells in response to position under normal conditions (blue line) and with grid cell depolarization (red line). All three cells are from the same network. **(a)**. Rate remapping. **(b)**. Place remapping. **(c)**. Global remapping. **(d)**. The effect of grid cell depolarization on the population activity of place cells. **(e)**. The histogram of bump scores of place cell activities across the environment.

Together, these simulation experiments demonstrate that our coupled network model can reproduce both the remapping induced by MEC inhibition and MEC depolarization, thereby providing a unified explanation for these seemingly contradictory experimental phenomena.

#### 5 Robustness test

In this section, we conduct simulation experiments to testify the robustness of our simulation results under the variation of some parameters.

#### 5.1 Impact of correlated noise

We conduct the same experiment as shown in Figure 3 and Figure 4 in the main text but with correlated multi-variate Gaussian noise, whose covariance matrix is,

$$\langle (R_i(x) - f_i)(r_j - f_j) \rangle = \sigma^2 \left[ \delta_{ij} + c(1 - \delta_{ij}) \right] \tag{81}$$

with $c = 0.15$, which falls within the range of experimentally observed noise correlations ($c = 0.1 - 0.2$) as reported by [15; 16]. These simulations confirmed that our main conclusions remain robust: (1) the network continues to integrate information in a Bayesian manner (see Figure S3a-b), and (2) even in the absence of positional information ($I_p = 0$), the network can still eliminate non-local errors caused by grid cell coding, enabling more robust spatial coding (see Figure S3c).

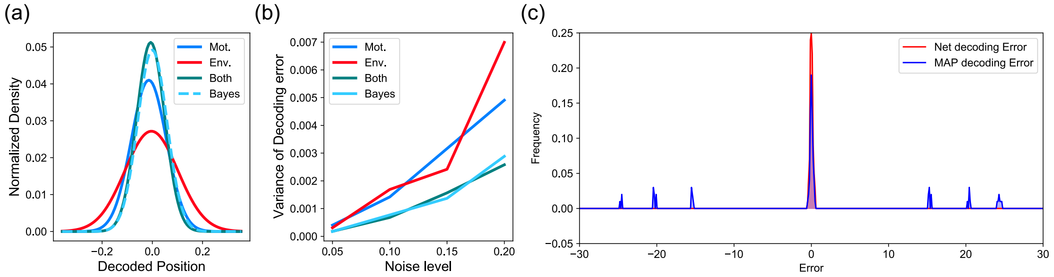


Figure S3: Simulation with Correlated Noise.

#### 5.2 Impact of Coupling Strength

We next examine whether variations in coupling strength (denoted as $J_{g_i,p}$ in Eq. (30)) affect the model's ability to integrate information. Specifically, we systematically varied the coupling strength and repeated the experiment shown in Figure 3 of the main text. The results, presented in Figure S4, demonstrate a robust alignment between the model's information integration and Bayesian integration across different coupling strengths.

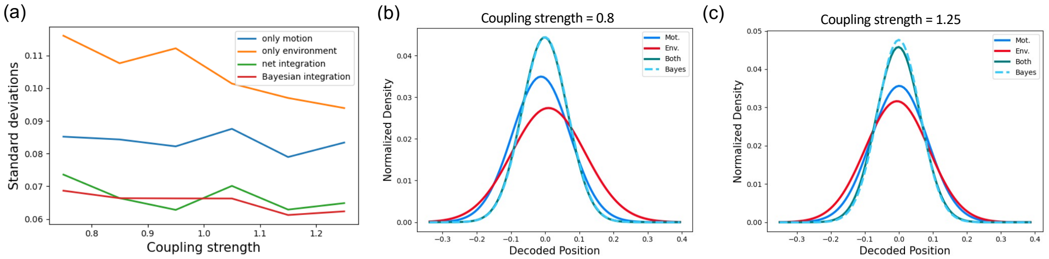


Figure S4: Impact of coupling strength on information integration. (a) Standard deviations of decoding results under different coupling strengths. (b) and (c) Examples of decoding result distributions for varying coupling strengths.

#### 5.3 Model Comparison

We compare our model with the "constrained range model" proposed by [1] (see Figure S5). Both models are capable of eliminating non-local errors when the decoding range is tightly constrained (Figure S5a). However, when the decoding range becomes larger, our model continues to eliminate non-local errors, whereas the "constrained range model" fails to do so (Figure S5b-c).

The key difference lies in how non-local errors are handled. Our model leverages recurrent currents within the place cell network to store historical information, enabling robust error correction without relying on range constraints. In contrast, the "constrained range model" reduces non-local errors by limiting the coding range, thereby sacrificing coding capacity. This distinction highlights the advantage of our model: the dynamics of the place cell network not only ensure robust spatial coding but also preserve coding efficiency.

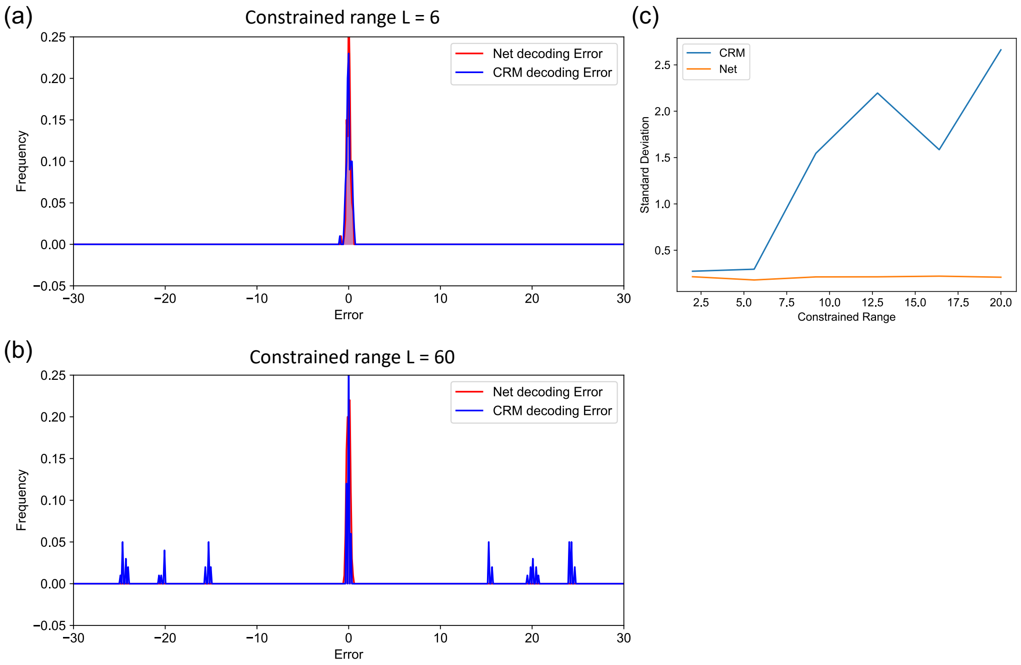


Figure S5: Comparison of our model with the constrained range model. (a) Both models successfully eliminate non-local errors when the decoding range is small. (b) and (c) When the decoding range is large, our model maintains error correction while the constrained range model fails.

#### 5.4 Capacity Dependence on Architectural Parameters

As demonstrated in Figure S6, we systematically evaluate the impact of two key architectural parameters on representational capacity: (1) The total number of neurons (Figure S6a): Although the representational capacity of both the coupled network and the only place cell network increases with the number of neurons, the coupled network consistently exhibits higher capacity than the only place cell network, confirming that grid cell integration enhances place coding efficiency; (2) The number of grid cell modules (Figure S6b): As the number of grid module ($M$) increases, the capacity of the coupled network increases monotonically. At low module numbers ($M < 6$), the coupled network's capacity is lower than that of the 1000 place cell network. This aligns with theoretical predictions, as grid cells require sufficient modular diversity to generate combinatorially efficient spatial encoding. When $M > 6$, the coupled network's capacity exceeds that of the 1000 place cell network.

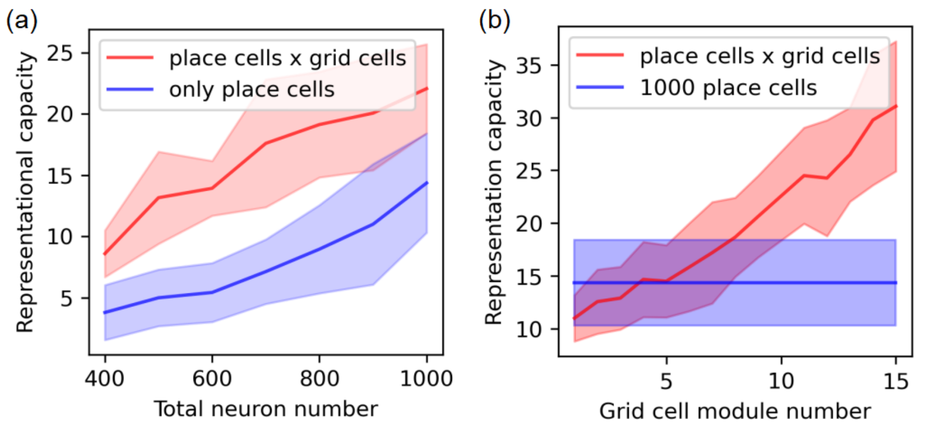


Figure S6: Representational capacity as a function of network architecture. (a) Representational capacity grows with the number of neurons for both the coupled network (80% place cells + 20% grid cells, red) and the only place cell network (blue). The coupled network achieves higher capacity at all population sizes due to the integration of grid cell dynamics. (b) Impact of modularity on representational capacity for a fixed population size (1,000 cells). The coupled network (800 place cells + 200 grid cells distributed across M modules, red) is contrasted with a homogeneous place cell network (1,000 place cells, blue). Data are shown as mean ± SD across 30 trials.

[14] K Wong, Si Wu, and Chi Fung. Tracking changing stimuli in continuous attractor neural networks. *Advances in Neural Information Processing Systems*, 21, 2008.

[15] Ehud Zohary, Michael N Shadlen, and William T Newsome. Correlated neuronal discharge rate and its implications for psychophysical performance. *Nature*, 370(6485):140–143, 1994.

[16] Michael N Shadlen, Kenneth H Britten, William T Newsome, and J Anthony Movshon. A computational analysis of the relationship between neuronal and behavioral responses to visual motion. *Journal of neuroscience*, 16(4):1486–1510, 1996.
